# Differential control of both cell cycle-regulated and quantitative histone mRNA expression by *Drosophila* Mute

**DOI:** 10.64898/2026.03.04.709556

**Authors:** Mark S. Geisler, James P. Kemp, Christina A. Hill, William F. Marzluff, Robert J. Duronio

## Abstract

Coupling histone gene expression to S phase of the cell cycle is essential for genome duplication and stability. Activation of Cyclin E/Cdk2 at the G1-S transition stimulates high-level expression of histone genes during S phase, but how histone genes are turned off at the end of S phase is not understood. Here we demonstrate that the essential *Drosophila* gene *mute* functions to repress inappropriate histone mRNA accumulation outside of S phase by counteracting Cyclin E/Cdk2-dependent phosphorylation of Mxc, which activates histone gene expression. Additionally, Mute plays contrasting roles in histone gene expression during S phase by promoting high levels of *H1*, *H2a* and *H2b* expression but not *H3* and *H4*. Although Mute is present only at replication-dependent histone genes, its loss leads to 801 differentially regulated genes, primarily those involved in muscle related processes in late-stage embryos. Thus, disruptions of histone gene expression control alters the transcriptome resulting in developmental defects.

## Introduction

During S phase of the cell division cycle millions of histone proteins are produced to assemble the nucleosomes needed to package newly replicated DNA into chromatin. For instance, each S phase a diploid human cell produces upward of 30 million nucleosomes comprised of 240 million histone proteins to package the new genome (Alberts B, Johnson A, Lewis J, 2002). This formidable biosynthetic task is accomplished by coordinating high level expression of multi-copy replication-dependent (RD) histone genes during S-phase, a feature of the cell cycle that is conserved from yeast to humans (Marzluff & Duronio, 2002; Osley, 1991). Changes in nucleosome occupancy, chromatin accessibility, and transcription from mis-regulation of histone production can result in chromosome instability, DNA damage sensitivity, cell cycle arrest, and defects in development (Alahmari et al., 2024; Amodeo et al., 2015; Celona et al., 2011; Chari et al., 2019; Clemente-Ruiz & Prado, 2009; Gunjan & Verreault, 2003; Han et al., 1987; Hauer et al., 2017; Shindo & Amodeo, 2021; Singh et al., 2010; Sullivan et al., 2001; Takayama & Toda, 2010). Reduced histone protein levels have been documented in aging human patients and senescent cells (Feser et al., 2010; Lowe et al., 2020; O’Sullivan et al., 2010; Song & Johnson, 2018), and increased RNA polymerase II (RNAPII) occupancy at RD-histone genes correlates with poor prognosis in multiple human cancer types (Henikoff et al., 2025). These observations highlight the important role regulation of histone production plays in genome integrity and human disease, and thus the need to understand mechanisms of histone gene regulation.

In metazoans, each RD histone protein is encoded by multiple genes that are sequestered in a phase separated nuclear biomolecular condensate called the Histone Locus Body (HLB) (Duronio & Marzluff, 2017; Geisler et al., 2023; Hur et al., 2020). In *Drosophila*, the RD-histone genes are present in a large array at a single locus containing ∼100 copies of each of the five RD histone genes (Bongartz & Schloissnig, 2019; Crain, Nevil, et al., 2024; McKay et al., 2015; Shukla et al., 2025). Transcription and processing of the unique RD histone mRNAs, which are not polyadenylated and instead end in a conserved stem loop, occurs within the HLB and the factors involved are conserved between *Drosophila* and humans (Kemp et al., 2021; Marzluff & Koreski, 2017; Skrajna et al., 2016; Yang et al., 2014). The *Drosophila* protein Mxc (NPAT in humans) is required for HLB formation, recruitment of other histone pre-mRNA processing factors, and for transcription of the RD histone genes (Tatomer et al., 2016; Terzo et al., 2015; White et al., 2011). Mxc/NPAT, along with other HLB components such as FLASH and RNAPII are present at HLBs throughout interphase of the cell cycle even though production of RD histone mRNAs is limited to S-phase (Armstrong et al., 2023; Hur et al., 2020; Kemp et al., 2025; Liu et al., 2006; Marmolejo et al., 2026; Terzo et al., 2015; White et al., 2007). Cyclin E/CDK2 phosphorylation of Mxc/NPAT activates RD-histone gene expression and accumulation of histone mRNA in the cytoplasm upon entry into S-phase (Armstrong et al., 2023; Kemp et al., 2025; Lanzotti et al., 2004; Ma et al., 2000; White et al., 2007; Ye et al., 2003; Zhao et al., 2000), including by stimulating elongation of promoter proximal paused RNA pol II (Kemp et al., 2025). Although many studies have investigated the activation of RD histone mRNA production at the beginning of S phase, relatively little is understood about how RD histone mRNA production is terminated as cells exit S phase.

Several proteins have been reported to repress RD histone mRNA accumulation (e.g. WGE, HERS, WEE1, HIRA, Abo, Histone H4, ATR) (Ahmad et al., 2026; Berloco et al., 2001; Hall et al., 2001; Ito et al., 2012; Mahajan et al., 2012; Marmolejo et al., 2026; Ozawa et al., 2016). Here we chose to focus on the *Drosophila* protein Mute (*muscle wasted*) (Bulchand et al., 2010), the orthologue of human protein Gon4L/YARP (Yang et al., 2014) because loss of Mute leads to increased accumulation of correctly processed H3 and H4 mRNA and there is strong evidence that Mute accumulates only in the HLB (Ahmad et al., 2026; Bulchand et al., 2010; Tatomer et al., 2016; White et al., 2011). Mute is a large, substantially disordered protein which contains roughly six predicted domains (Fleming et al., 2025; Jumper et al., 2021), three of which are homologous to predicted protein-protein binding domains: two Paired-Amphipathic Helix (PAH) domains, similar to those found in the Sin3a repressive hub protein (Bugge et al., 2021; Sahu et al., 2008), and a C-terminal SANT/Myb domain which interacts with Mxc *in vitro* (Yang et al., 2014) (Fig 1A). These domains and interactions are conserved between human Gon4L and NPAT (Bucholc et al., 2020). The Gon4L PAH domains interact with CRAMP1 (Cramped in *Drosophila*), which is required for *H1* gene expression but not for core RD histone gene expression (Bodner et al., 2026; Gibert & Karch, 2011; Ingham et al., 2025; Matthews et al., 2025; Yamamoto et al., 1997). Although analyses of *mute* mutant phenotypes indicate a role for Mute in repression of RD histone gene expression, if and how Mute contributes to the cell-cycle-dependent regulation of RD histone genes, and whether it affects all histone genes similarly, has not been determined. Loss of Mute causes lethality in late-stage embryos (Bulchand et al., 2010), but whether Mute is required for cell proliferation is not known.

**Figure 1.**
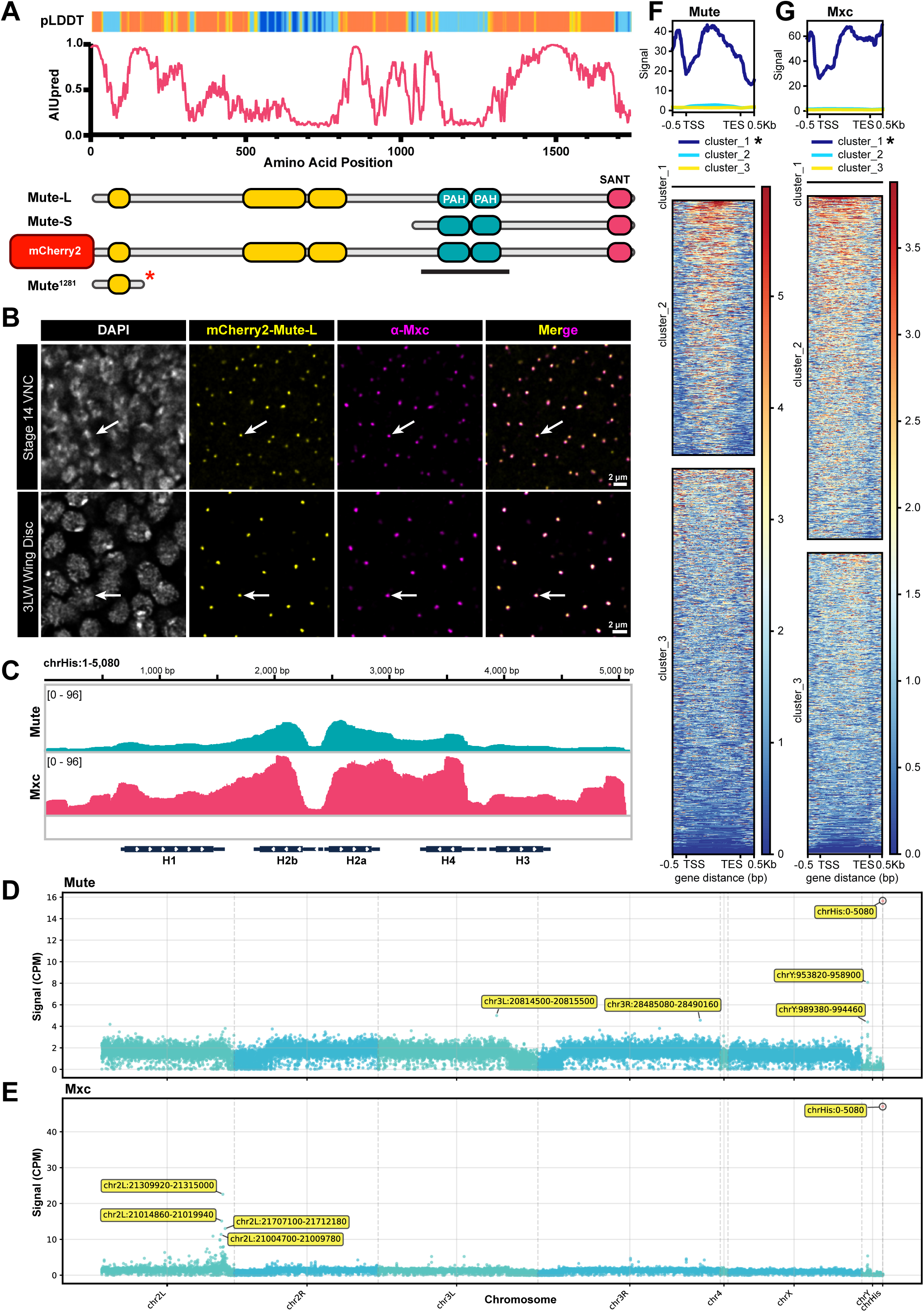
Mute localizes only to RD histone genes **A)** Schematic of the two protein isoforms encoded by the *mute* locus, the mCherry2 N-terminally tagged Mute-L, and *mute^1281^* alleles overlayed with AIUpred disorder prediction (1 is highest probability of a residue in disordered region) and per residue pLDDT from AlphaFold (dark blue is highest structural confidence) (Erdős & Dosztányi, 2024; Jumper et al., 2021). Yellow areas are unassigned but predicted protein domains. Black bar indicates epitope used to generate α-Mute-LS antibody in Bulchand et al. 2010 (Bulchand et al., 2010) **B)** Stage 14 embryonic ventral nerve cord and third larval instar wing imaginal discs expressing mCherry2-Mute-L (yellow) and stained with antibodies against Mxc (magenta) and DAPI (white). Note the co-localization of Mute and Mxc and the lack of detectable nuclear signal outside of the HLB (arrows). Scale bar = 2μm. **C)** IGV screenshot of “per histone gene copy” (HGC) normalized CUT&RUN signal at the 5080nt histone gene repeat using anti-Mute and anti-Mxc antibodies and third instar larval wing discs. **D,E)** Manhattan plots of Mute (D) or Mxc (E) signal (CPM) in 5080nt bins across the entire *Drosophila* genome, with the histone gene repeat unit entered as a distinct “chromosome” (chrHis). The top five bins are labeled, including the HGC signal at chrHis (red dot at far right). **F,G)** Metagene plot and heatmap of summed HGC normalized Mute (F) and Mxc (G) signal across all *Drosophila* genes (17,266 genes) +/- 0.5 kb upstream of the TSS and downstream of the TES. Genes were clustered using unsupervised hierarchical clustering for n = 3 bins. In both Mute and Mxc, the histone genes clustered by themselves in cluster 1 (asterisk).

Here we show that Mute localizes only to the HLB and RD histone genes and is required for restricting the expression of all five RD histone genes to S phase. Interestingly, we show using multiple methods that Mute regulates the expression of H3 and H4 differently than H1, H2a and H2b, highlighting that a single regulator within a single biomolecular condensate can mediate differential gene regulation. Our data suggests a model whereby Mute controls the cell cycle-coupled expression of all five RD histone genes, terminating expression at the end of S phase by opposing Cyclin E/Cdk2 phosphorylation of Mxc, while also controlling the quantity of H1, H2a and H2b mRNA expressed during S phase. This regulation is critical for development, as loss of Mute disrupts the transcriptome during embryonic development and blocks cell proliferation in larvae.

## Results

### Mute localizes only to the RD histone genes and Histone Locus Body

Mute was previously shown to localize to the HLB via immunofluorescence using antibodies raised against both the N- and C-terminus, the latter of which detects both the long and short isoform of Mute (Bulchand et al., 2010). To extend this finding, we used CRISPR directed homologous repair to generate an endogenously tagged Mute protein by fusing mCherry2 to the N-terminus of the long isoform of Mute (Mute-L) (Fig. 1A). We found that mCherry2-Mute-L is present in the HLB as it colocalizes with antibodies against Mxc in both late-stage embryos and third instar wing imaginal discs, which are a population of asynchronously cycling epithelial cells that are precursors of the adult wing and thorax (Fig. 1B, arrows). Like the HLB factors Mxc, FLASH, and U7snRNP, Mute is present in the HLBs of all wing disc cells, indicating that its localization to the HLB is not cell cycle dependent. Moreover, we cannot detect appreciable accumulation of Mute outside of the HLB. CUT&RUN analyses of wing imaginal discs using anti-Mute and anti-Mxc antibodies revealed strong binding of both Mute and Mxc to RD histone genes (Fig. 1C) as also seen with ChIP-seq (Hodkinson et al., 2024) and CUT&TAG (Ahmad et al., 2026).

We next assessed whether Mute and Mxc bound loci other than the RD histone locus. To accommodate mapping CUT&RUN data resulting from the ∼100 histone gene repeat cluster (Bongartz & Schloissnig, 2019), paired end reads were aligned to a custom dm6 genome where the partially assembled RD histone locus (*HisC*; chr2L:21,400,839-21,573,418) was removed and replaced with a single copy of the 5080bp histone gene repeat as its own chromosome (chrHis) (Fig. 1C). chrHis reads in Counts Per Million (CPM) normalized BigWigs were divided by 100 to achieve “per histone gene copy” (HGC) signal within chrHis to compare to the rest of the genome. We subdivided the genome into 5080bp non-overlapping bins and summed the Mute and Mxc signal within each bin. For both proteins, chrHis had significantly higher HGC signal than any other genomic bin (Fig. 1D,E). The other four bins with the highest Mxc signal occur within a few hundred kilobases on either side of *HisC* on chr 2L, which could conceivably be in close proximity to the HLB and thus Mxc (Fig S1A). For Mute, the next highest four bins include two intergenic, repeat heavy chrY regions and two bins with much lower signal on either arm of chr3 (L and R) Fig S1B,C). We also summed Mute and Mxc signal across all annotated genes in the *Drosophila* genome then performed unsupervised hierarchical clustering (Fig. 1F,G). In both cases, all five RD-histone genes clustered by themselves, showing high signal across the genes (Fig. 1F,G, top). All other genes showed low to no signal and did not appear to cluster in any meaningful way. We conclude from these data that Mute and Mxc strongly associate only with RD histone genes, suggesting that the primary, and perhaps only, function of Mute occurs at the histone locus.

### Loss of Mute increases the number of cells expressing core RD histone genes in the *Drosophila* embryonic ventral nerve cord

Hemizygosity for the *mute^1281^*null allele, which contains a nonsense mutation near the beginning of the Mute-L open reading frame (Fig. 1A), results in embryonic lethality and an increase in correctly processed histone H3 and H4 mRNA in late stage mutant embryos (Bulchand et al., 2010). The latter result was obtained by Northern blotting of RNA extracted from late-stage embryos and thus could not assess RD histone gene expression at the single cell level or with respect to cell cycle progression. Using RNA-FISH probes that hybridize to the coding regions of the four core RD histone genes (*H2A*, *H2B*, *H3*, and *H4*) (Fig. 2A)(Kemp et al., 2025), we assessed RD histone mRNA levels throughout development of homozygous *mute^1281^* embryos and their heterozygous siblings. We noticed a striking phenotype in the Ventral Nerve Cord (VNC) of stage 14 embryos in which more cells contain histone mRNA in the *mute^1281^* genotype compared to heterozygous siblings (Fig. 2B,C). At higher magnification we detected FISH signal that colocalizes with Mxc, representing nascent transcription of the RD histone genes (Fig. 2D, arrows). Nascent transcription can be detected in the VNC cells containing histone mRNA in the *mute^1281^* null embryos, suggesting that Mute regulates transcription of RD histone genes rather than RD histone mRNA stability. Antibodies raised against the C-terminus of Mute do not stain HLBs of stage 14 *mute^1281^* embryos (Fig. 2E) nor mute deficiency embryos that lack the entire *mute* locus (Fig. S2A), indicating that any maternal load of Mute has been degraded by this point in embryogenesis. Some persistence of maternally loaded Mute is likely why the RD histone mRNA phenotype does not appear until stage 14. Loss of Mute does not disrupt HLB formation as assessed by staining *mute^1281^* HLBs with antibodies recognizing Mxc or FLASH (Fig. 2D,E).

**Figure 2.**
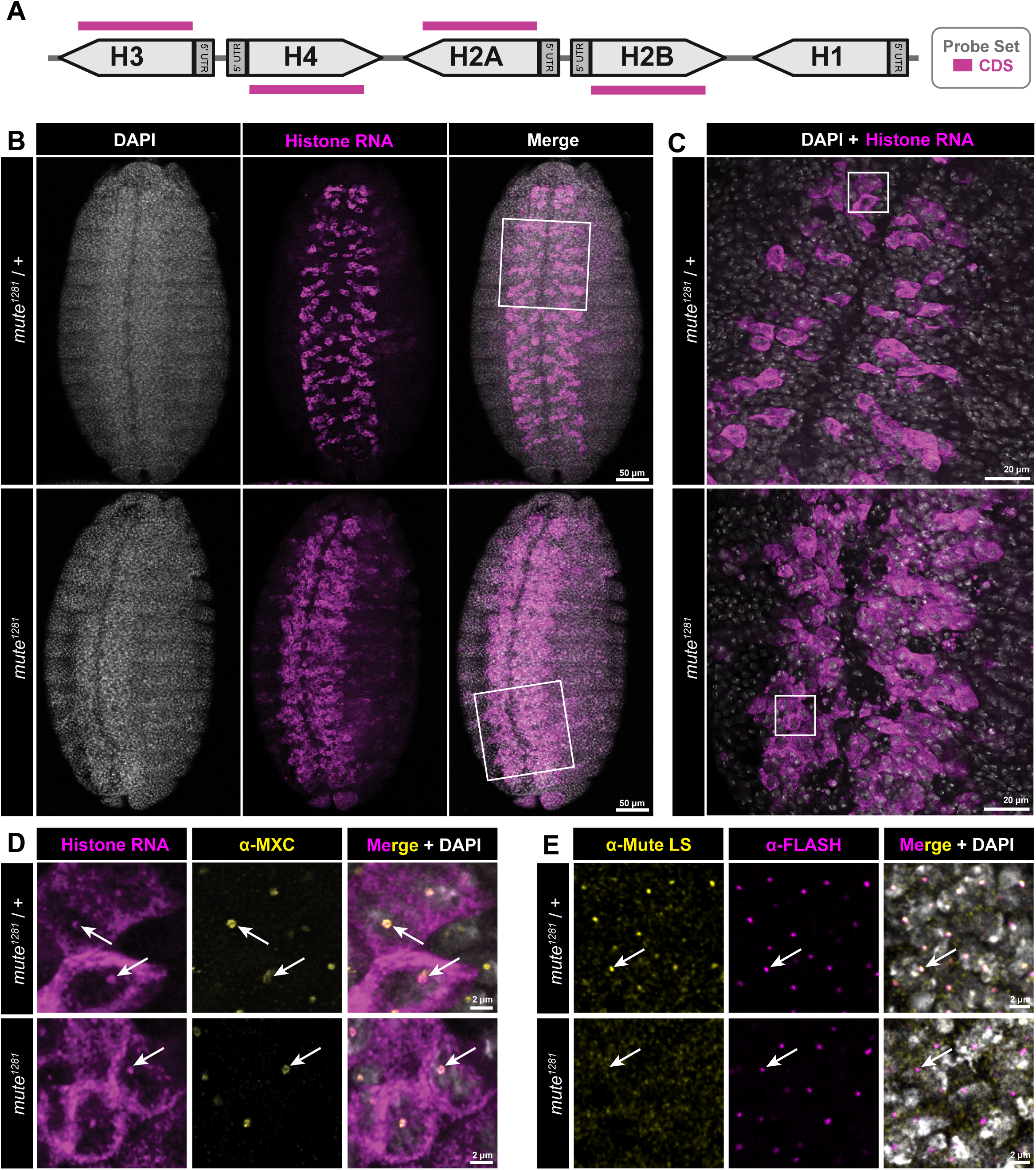
Ectopic core RD histone gene expression in *mute* mutant embryonic VNC **A)** Schematic of a single RD histone gene unit and the location of the fluorescently labeled oligonucleotide probe set (CDS) used to detect transcription and accumulation of core RD histone mRNA. **B)** Ventral view of stage 14 *mute^1281^* null and heterozygous sibling control embryos hybridized with the CDS probe (magenta) and stained with DAPI (white) and anti-Mxc antibodies (yellow, panel D). White boxes indicate regions imaged in higher magnification in panel C. Scale bar = 50μm. **C)** Higher magnification view of the VNC showing ectopic expression of core RD histone genes. White boxes indicate corresponding regions shown in panel D. Scale bar = 20μm. **D)** Highest magnification view of *mute^1281^* null and heterozygous sibling control embryos indicating accumulation of nascent core RD histone transcripts within the HLB (arrows). Scale bar = 2μm. **E)** *mute^1281^* null and heterozygous sibling control embryos stained with anti-Mute (yellow) and anti-FLASH (magenta) antibodies. Note the absence of Mute accumulation in the HLB in the *mute^1281^* null genotype (arrows). Scale bar = 2μm.

The embryonic VNC has highly regulated patterns of cell division and differentiation (Bier et al., 1992; Campos-Ortega, 1999; Vaessin et al., 1991). Most embryonic neuroblasts (NBs), the neural stem cells of *Drosophila,* undergo a type I lineage of self-renewal, asymmetrically dividing into another NB and a Ganglion Mother Cell (GMC) daughter cell (Campos-Ortega, 1999) (Fig. S2B). This process is governed by numerous neural transcription factors (TF), including the NB-specific factor Deadpan (Dpn), which is expressed in NBs and transcriptionally silenced in GMCs by the neural differentiation factor Prospero (Pros) (Bier et al., 1992; Vaessin et al., 1991). We asked if the loss of Mute, and potentially the misexpression of the RD histone genes, disrupted these patterns and lineages. We co-stained wildtype *OregonR* and *mute^1281^* stage 14 embryos with antibodies against Pros and Dpn (Fig. S2C,D). The number of cells staining for Dpn was two-fold higher in *mute^1281^* stage 14 embryos compared to control (Fig. S2C,E) and there was a smaller but significant increase in the number of cells staining for Pros (Fig S2F). In *mute^1281^* embryos, 50% of cells positive for Dpn also contained Pros, up from 27% in wildtype (Fig. S2D, bottom, yellow circle; S2E). In wild type embryos we can identify many Pros-positive cells with low Dpn that lack RD histone mRNA (Fig. S2D, top, circle; S2E,F). By contrast, almost all Pros-positive GMCs in *mute^1281^* mutant embryos express RD histone mRNA (Fig. S2D, bottom, white circle; S2F,G). Thus Mute, and likely correct regulation of histone gene expression, is necessary for normal development of the embryonic VNC. These results are consistent with previous work demonstrating that Ap4/FMRFa neurons are absent in *mute^1281^* embryos (Bivik et al., 2015).

### Mute is required to restrict core RD-histone gene expression to S-phase

Dividing cells of the VNC undergo a condensed S-G_2_-M cell cycle with cells exiting into a G_1_ arrest after the cessation of proliferation and subsequent differentiation (de Nooij et al., 1996; Fichelson et al., 2005; Lane et al., 1996; Weigmann & Lehner, 1995). To determine if the increased numbers of VNC cells expressing RD histone genes in *mute^1281^* embryos were in S phase, we combined 5-ethynyl-2’-deoxyuridine (EdU) labeling to mark DNA replication with RNA-FISH to detect RD histone RNA (Fig. 3A). We quantified the number of cells in the VNCs of *OregonR*, *mute^1281^* heterozygous, and *mute^1281^* homozygous mutant embryos that labeled with EdU and accumulated cytoplasmic histone RNA (Fig. 3B, top 2 rows, 3C-D), or that accumulated RD histone mRNA but did not label with EdU (Fig. 3B, bottom, 3C-D). As expected, in both *OregonR* and the *mute^1281^* heterozygous siblings there was substantial cooccurrence of EdU labeling and cytoplasmic, RD histone RNA (Fig. 3C,D). By contrast, in *mute^1281^* embryos we observed an over 7-fold increase relative to control in the number of cells which contained cytoplasmic RD-histone mRNA but did not label with EdU (Fig. 3C,D). In these cells we also detected RD-histone RNA-FISH signal at the HLB, indicating transcription of RD histone genes occurring outside of S-phase. There is not a significant difference in the total number of S phase cells between the controls and *mute^1281^* mutants, indicating that the *mute^1281^* mutation does not substantially impede progression through S phase at this early stage of development (Fig. 3D). We conclude from these data that Mute is required to restrict RD histone gene expression to S phase and that loss of Mute results in aberrant RD histone gene expression outside of S phase.

**Figure 3.**
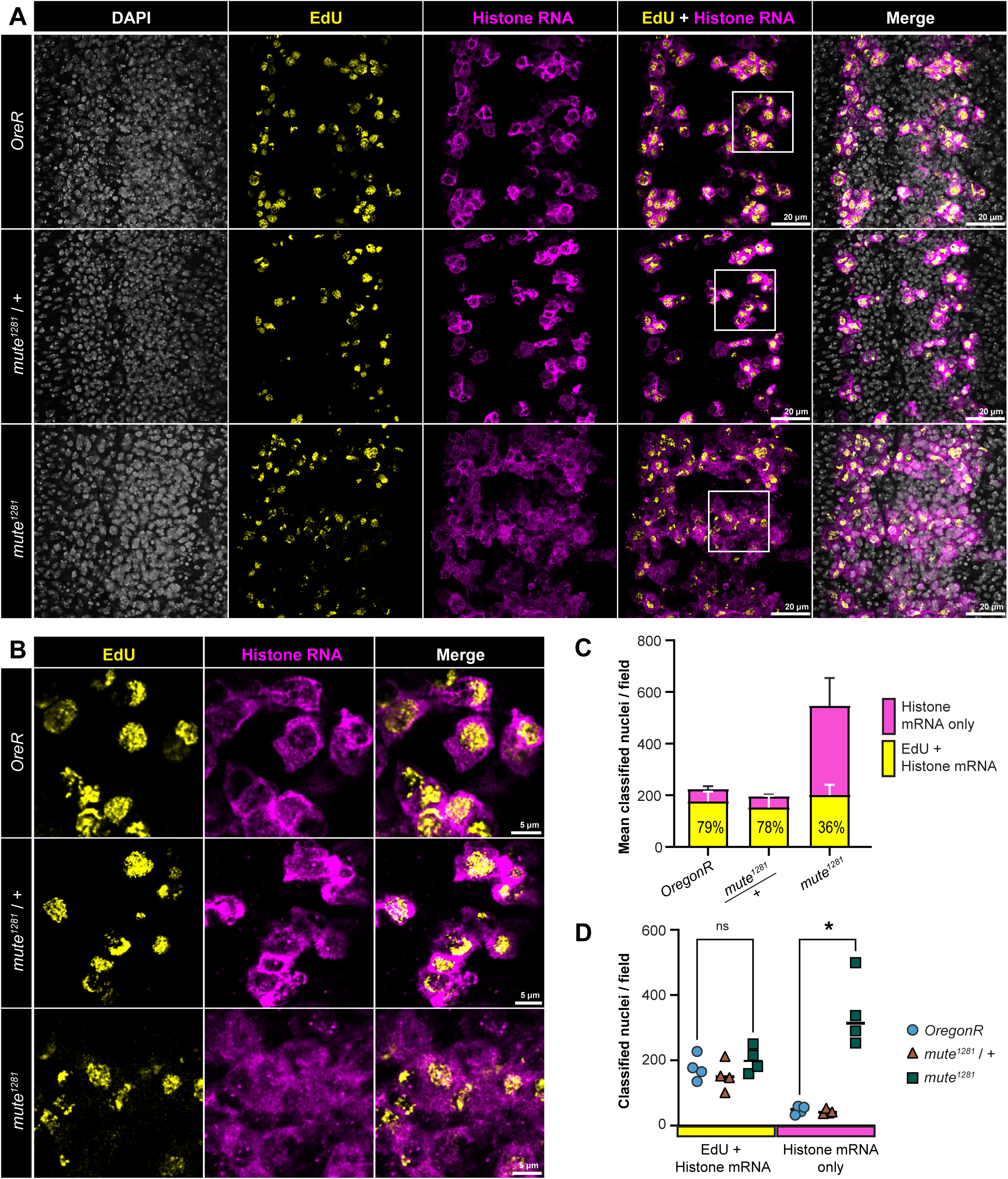
Core RD histone mRNA accumulates outside of S phase in the *mute* mutant VNC **A)** VNC of wild type *OregonR*, *mute^1281^* null, and *mute^1281^* heterozygous sibling embryos pulse labeled with EdU (yellow) and hybridized with the CDS probe (magenta). White boxes indicate corresponding regions shown in panel B. Scale bar = 2μm. **B)** Higher magnification view of embryos from panel A. Scale bar = 5μm. **C, D)** Quantification of proportion of embryonic VNC cells containing core RD histone genes without incorporating EdU in the indicated genotypes. Error bars are +/- standard deviation of n = 4 replicates. Significance determined by unpaired T-test with FDR; * = pval < 0.001; ns = no significance.

### Mute couples expression of all RD histone genes to S phase but differentially regulates level of RD histone genes during S phase

Current models suggest that genes encoding the nucleosome core are regulated as a unit, since stoichiometric production is necessary to promote the assembly of functional nucleosomes. To test this model, we used differently colored RNA-FISH probe sets that hybridize to single histone RNAs (Fig. S3A). We determined the expression patterns and relative levels of H1, H2a, and H3 mRNA in individual cells of the VNC. The patterns of expression of these individual genes were different than wild type and the same as we detected with the CDS probe recognizing all 4 core histone genes (Fig. 4A). To quantify the results, pixel intensities of a 1.8 µm deep z-stack centered on the VNC were summed and then binned to generate histograms of pixel intensities across the 100 µm^2^ images (Fig. S3C). These histograms show a rightward shift in *mute^1281^* embryos, meaning there is more signal throughout the entire area of the image, which matches what is apparent by eye. This reduction in skewness (shift toward a normal distribution) in *mute^1281^* is significant for all four probe sets (Fig. 4B) and consistent with an increase in the number of cells accumulating RD histone mRNA because of gene expression outside of S-phase. RNA-FISH probes hybridizing to either H1 or H2a RNA revealed lower maximal signal intensity in *mute^1281^* VNC compared to wildtype, an effect not seen with the H3 and CDS probes (Fig. 4A). The reduced maximal H1 and H2a but not H3 expression is apparent when comparing the bin value of the 90^th^ percentile (meaning 90% of all pixels are less than that value) (Fig. 4C, D-E dashed lines), or when plotting pixel intensity histograms on a log scale for H1 (Fig. 4E), H2a (Fig. S3B), and H3 (Fig. 4D). These data reveal that loss of Mute differentially affects the maximal expression of some RD histone genes but is required to restrict expression of all RD histone genes to S phase (Fig. 4L).

**Figure 4.**
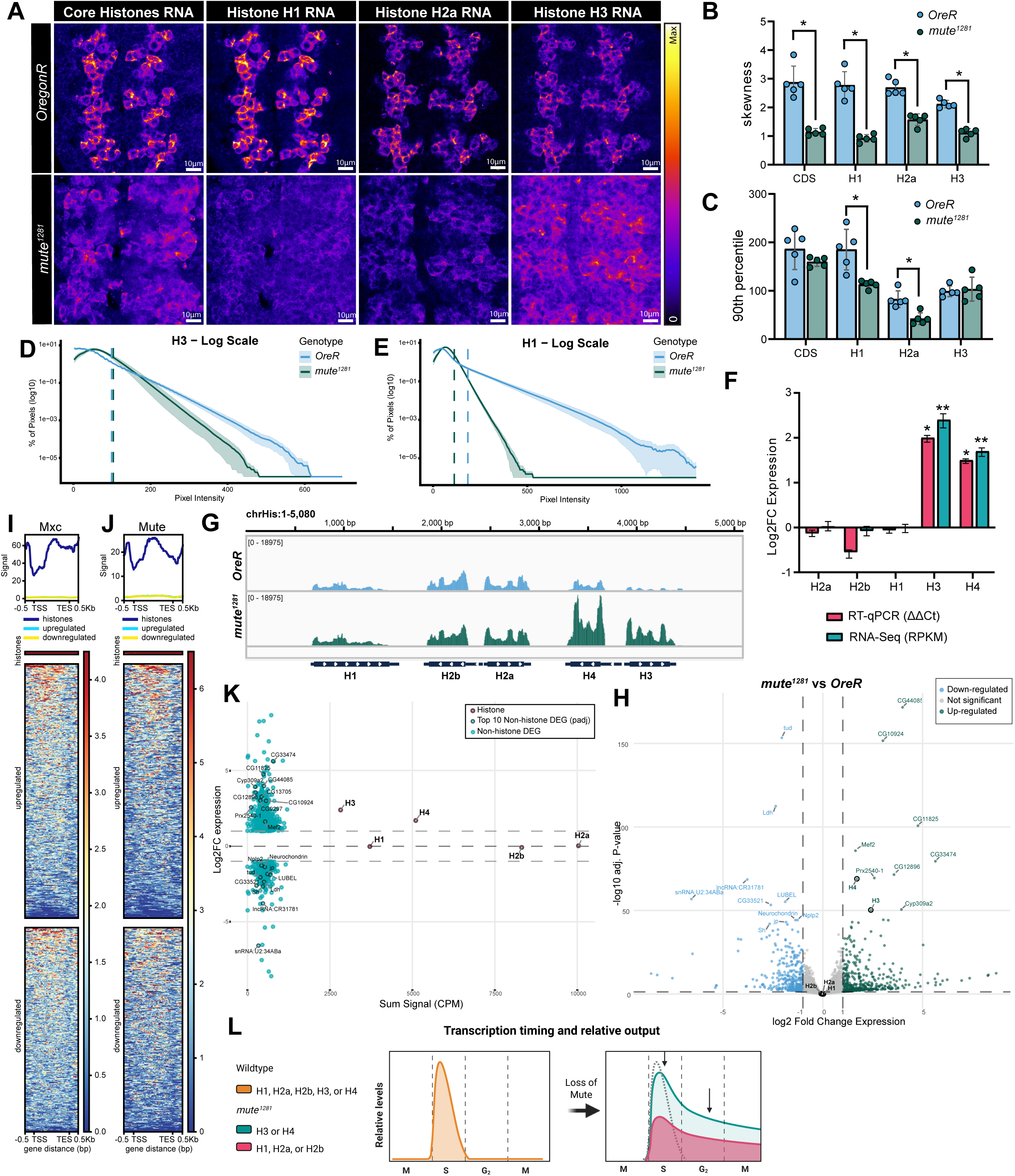
Mute restricts all histone gene expression to S-phase but differentially affects maximum expression of H1, H2a and H2b versus H3 and H4. **A)** Intensity Sum projections of the VNC of *OreR* and *mute^1281^* embryos hybridized with either both the CDS and Histone H1 probes or both the Histone H2a and Histone H3 probes. Images for each probe set are contrast adjusted identically across genotypes with the gradient LUT showing relative intensity of RNA levels. For visualization only, max pixel values for both genotypes are 615 (CDS), 615 (H1), 375 (H2a), and 450 (H3) for the different probe sets. Scale bar = 10μm. **B)** Bar graph showing change in skewness (distribution asymmetry) between *OreR* (blue) and *mute^1281^* genotypes of binned pixel intensity sum measurements across 5 VNCs per genotype per probe set. Decrease in skewness represents a shift to more normal distribution of data. Error bars = SD; Mann-Whitney U test for n =5 replicates per probe set per genotype. * = adjusted pval (padj) < 0.05 **C**) Bar graph of 90^th^ percentile bin value for each probe set across genotypes. Test same as B, * = padj < 0.05 **D,E)** Log_10_ transformed histogram of mean H3 (D) or H1(E) probe set counts per pixel value in each genotype. Solid line represents mean of n=5 with SEM shaded bars. Dashed vertical lines represent average 90^th^ percentile start for each genotype. **F)** Bar graph of Log_2_ fold change (Log_2_FC) in expression for all five RD-histone genes measured by RT-qPCR (ΔΔC^t^) and RNA-Seq (RPKM) in *mute^1281^*embryos compared to *OregonR*. Both techniques show significant Log_2_FC only for H3 and H4. *=p< 0.005 (two-tailed t-test with BH FDR), **=padj < 1e-50; Error bars = lfcSE; n = 3 **G)** IGV browser shot of RPKM normalized signal of combined strands at chrHis in *OregonR* (blue) and *mute^1281^* (green). Note increase in signal only at H3 and H4. **H)** Volcano plot of Log_2_FC expression vs -log_10_ adjusted P-value of all genes in *mute^1281^* vs *OregonR* embryos. Genes with Log_2_FC greater than +/- 1 and -log_10_ adj. p-value greater than 1.3 (p-val < 0.05) are differentially down-(blue) or up-regulated (green) genes. The top 10 most significant up and down DEGs are labeled. Unchanged genes, including H1, H2a, and H2b, are colored grey. **I,J)** Metagene plot and heatmap of summed HGC normalized Mxc (I) and Mute (J) signal across all RD-histone genes (5 genes, dark blue), non-histone upregulated genes (448 genes, light blue) and downregulated genes (351 genes, yellow) normalized to 2kb +/- 0.5 kb of the TSS and TES. **K)** MA plots of summed Mute signal (CPM) vs Log_2_FC expression at all RD-histone genes and DEGs (green). Top 10 up and down DEGs by adj. p-val are labeled. **L)** Model of Mute’s multiple roles in regulating histone gene expression. In individual cells, Mute restricts expression of all five RD-histone genes to S-phase and promotes full expression of H1, H2a, and H2b, but not H3 and H4. Consequently, when molecularly measuring mRNA in whole embryos, H3 and H4 mRNA accumulation (green) appears elevated whereas H1, H2a, and H2b do not (red).

To measure the bulk RNA level for each RD histone gene molecularly, we extracted RNA from 50 whole wild type or *mute^1281^* stage 14 embryos and performed both reverse transcription quantitative PCR (RT-qPCR) and paired end RNA sequencing (RNA-seq). Using both assays, we saw no significant change in Histone H1, H2a, and H2b RNA but a >1.5 Log_2_ fold change in both H3 and H4 in *mute^1281^* embryos (Fig. 4F). This lack of change in total H2a, H2b, and H1 mRNA accumulation at the bulk level may be explained by lowered max transcriptional output (Fig. 4E) that extends throughout the cell cycle (Fig. 4A) leading to a similar total amount of mRNA produced per embryo (shaded area under the curve in Fig. 4L). The max transcriptional output H3 and H4 in individual cells is not significantly different from wildtype (Fig. 4D) and when extended throughout the cell cycle results in higher accumulation of H3 and H4 mRNA at the bulk level (Fig. 4F,G). We detected no change in expression of any RD histone gene by RT-qPCR of heterozygous siblings (Fig. S3E) or of any replication-independent histone variant gene (Fig. S3F).

### Loss of Mute indirectly leads to differential expression of muscle related genes

To determine the consequence of Mute loss on the transcriptome we performed differential expression analysis with the RNAseq data using DEseq2 (Love et al., 2014). We found 801 significant differentially expressed genes (DEGs), consisting of 351 down regulated and 450 upregulated including H3 and H4, but not the other three RD-histone genes (Fig. 4H). Using PANGEA’s gene set enrichment analysis (GSEA) built on direct Gene Ontology (GO) Biological Processes (Hu et al., 2025), the most significantly enriched Gene Sets among these DEGs are associated with muscle system processes, muscle contraction, and transmembrane transport (Fig. S4D). GSEA using a gene set built on preferred tissue from modEncode showed the highest and most significant enrichment in muscle and digestive system tissues (Fig. S4E). These changes in expression of genes related to muscle processes and cells may explain the muscle wasted phenotype previously seen in later stage *mute^1281^* embryos (Bulchand et al., 2010).

To determine whether these muscle defects are direct or secondary effects from misregulation of histone gene expression upon loss of Mute, we evaluated both Mxc and Mute HGC CUT&RUN signal at RD histone genes along with the up and down DEGs in *mute^1281^* embryos (Fig. 4I,J). Both heatmaps and metagene plots for these three categories show no enrichment of either Mxc or Mute at both up and down DEGs. Since the histone vs differential clusters used in the metagene plots contained very different numbers of genes (5 vs. 400), we also plotted the Log_2_ fold change in expression against sum of HGC Mute signal for each differentially regulated gene and all five RD histone genes (Fig 4K). Other than *H3* and *H4*, no differentially expressed gene, either up or down, had nearly the same Mute signal as any of the RD-histone genes. There also is no correlation between Mute signal, however low, and relative changes in expression. We conclude that DEGs and the muscle and other developmental defects in *mute^1281^* embryos are not a direct result of Mute binding to non-histone genes, and thus likely result from pleiotropic effects due to misregulation of RD histone gene expression.

### Without Mute core RD histone genes are inappropriately expressed into G2/M

We next examined more thoroughly the uncoupling of RD histone gene expression from S phase we observed in the *mute^1281^* embryonic VNC. To determine cell cycle stage of *mute^1281^* cells we utilized a fly-FUCCI system that ubiquitously expresses a fragment of the Cyclin B protein fused to both a nuclear localization signal (NLS) and a fluorescent protein (RFP) (Zielke et al., 2014). Cyclin B-RFP is absent during G1 phase, begins to accumulate during early S phase and increases in expression throughout the remainder of interphase, peaking as cells enter mitosis when APC/C coordinates its degradation at the initiation of anaphase (Zielke et al., 2014)(Fig. 5C). We observed in control embryos that G2 cells with the highest levels of Cyclin B-RFP do not accumulate RD histone mRNA (Fig. 5A, top, arrow), whereas cells with lower levels of Cyclin B-RFP and thus in S phase have cytoplasmic histone RNA (Fig. 5A, top, double arrow). By contrast, VNC cells in *Mute^1281^* embryos with any detectable level of Cyclin B-RFP also contain cytoplasmic RD histone RNA (Fig. 5A, bottom, arrow; 5E). The total number of VNC cells in S phase or expressing Cyclin B-RFP are not statistically different between control and *Mute^1281^* embryos (Fig. 5D). These data are consistent with aberrant expression of RD histone genes in G2 upon loss of Mute (Fig. 5C).

**Figure 5.**
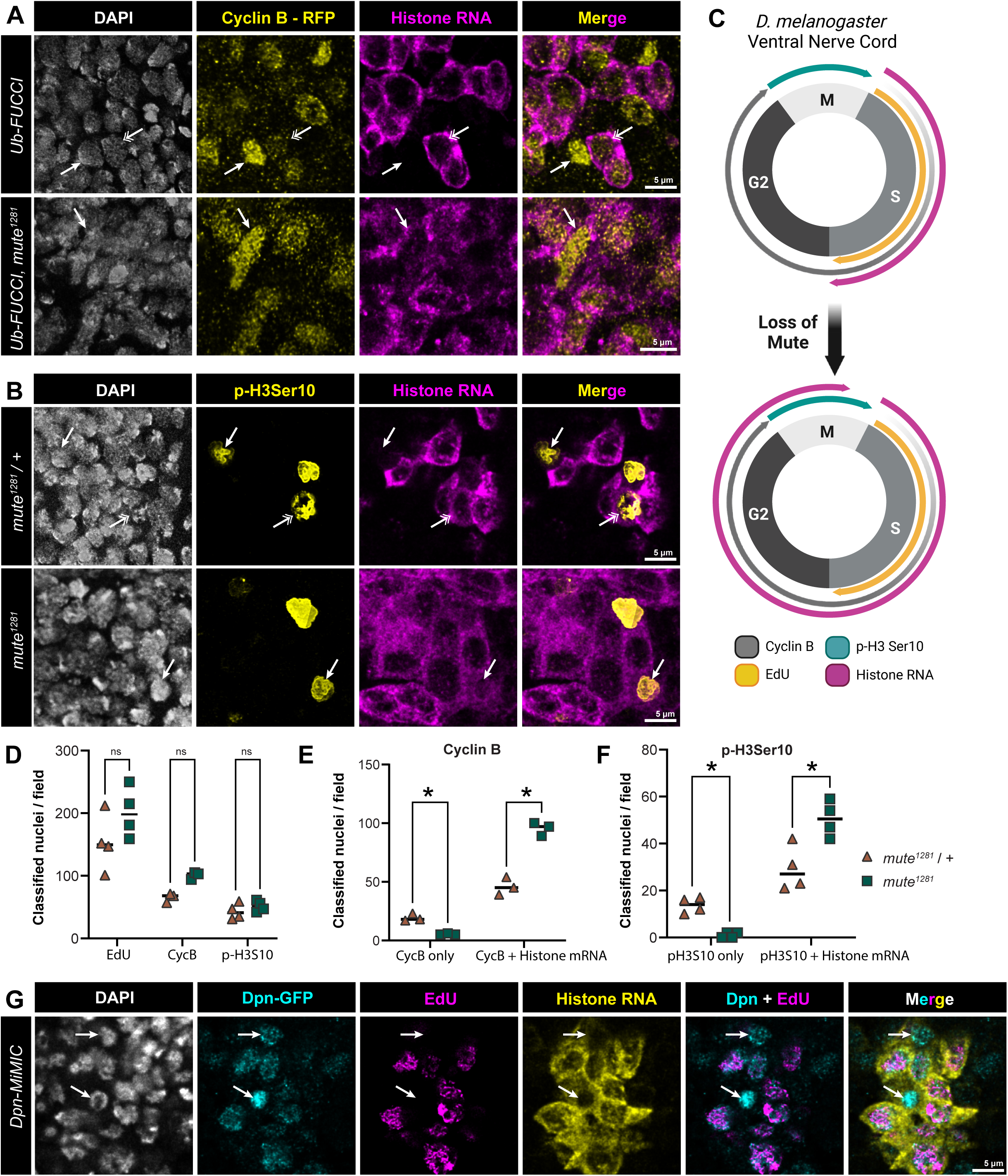
*mute* mutant embryonic VNC cells inappropriately express core RD histone mRNAs in G2 phase **A)** VNC of control and *mute^1281^* null ub-FUCCI embryos expressing Cyclin B-RFP stained with α-RFP antibody (yellow) and hybridized with the CDS probe (magenta). Single arrows indicate G2 cells expressing high levels of Cyclin B-RFP, double arrows indicate s-phase cells with lower levels of Cyclin B-RFP and cytoplasmic histone RNA. Scale bar = 5μm. **B)** VNC of *mute^1281^* null and heterozygous sibling control embryos hybridized with the CDS probe (magenta) and stained with antibodies against phospho-Ser10 of H3 (p-H3Ser10) to indicate mitotic chromosomes (yellow). Arrows indicate mitotic cells, which express RD histone mRNA more often in the *mute* mutant than in control (double arrow). Scale bar = 5μm. **C)** Schematic of expression of cell cycle markers and histone mRNA in wild type and *mute* mutant VNC cells. **D-F)** Quantification of the data shown in panels B and C. Paired t-test with FDR (D,E) or unpaired t-test with welch correction, FDR used. n = 4 (EdU, pH3) or 3 (CycB); * = pval < 0.05; ns = no significance **G)** VNC of Deadpan-GFP-expressing neuroblasts (cyan) in stage 14 embryos pulse labeled with EdU (magenta) and stained with DAPI (white) and hybridized with the CDS probe (yellow). Arrows show G2 neuroblasts (Deadpan positive, EdU negative) with histone mRNA. Scale bar = 5μm.

To determine if expression of RD histone genes is extending into mitosis upon loss of Mute, we stained embryos with antibodies recognizing H3 phospho-Serine 10 (p-H3S10), a histone post-translational modification (PTM) which marks condensing chromosomes during mitosis (Sauvé et al., 1999; Wei et al., 1999). In *mute^1281^* heterozygous control sibling embryos we detect cells that stain for p-H3S10 and do not have any discernable RD histone mRNA accumulation (Fig. 5B, top, closed arrow). In *mute^1281^* embryos, all cells positive for p-H3S10 also contain RD histone mRNA (Fig. 5B, bottom, closed arrow, Fig. 5F). We did not observe p-H3S10 positive cells that lack RD histone mRNA in *mute^1281^* embryos (Fig. 5F). As with Cyclin B-RFP, the total number of p-H3S10 positive cells is indistinguishable between control and *mute^1281^* embryos (Fig 5D). These data indicate inappropriate continued accumulation of RD histone mRNAs in mitosis in the absence of Mute.

Interestingly, in control embryos we observe some cells positive for p-H3S10 that accumulate cytoplasmic RD histone RNA (Fig. 5B, top, double arrow, Fig. 5F). These cells are likely neuroblasts (NB), which express Cyclin E continuously throughout the cell cycle (Betschinger et al., 2006; Weigmann & Lehner, 1995) thereby activating RD histone gene expression outside of S phase. To determine whether NBs normally express RD histone genes outside of S phase, we utilized GFP-tagged Deadpan (Dpn), a marker of NBs. We performed EdU labeling with RNA-FISH against the core RD-histone genes in stage 14 wild type embryos expressing GFP-Dpn and observed a population of GFP-Dpn-positive NBs which did not label with EdU and contained cytoplasmic RD histone RNA (Fig. 5G, arrows). This result indicates that cycling NBs can express RD-histone genes outside of S-phase, probably due to their continuous expression of Cyclin E throughout the S-G2-M cell cycle (Betschinger et al., 2006). This property of NBs likely explains the 20% of wild type VNC cells in Fig. 3C that express RD histone mRNA and do not label with EdU.

### Cyclin E/Cdk2 phosphorylation of Mxc is negatively regulated by Mute

Activation of Cyclin E/Cdk2 at the G1-S transition results in initiation of DNA replication and phosphorylation of Mxc, which induces the synthesis of RD histone RNA by stimulating RNA pol II elongation on RD histone genes (Kemp et al., 2025) and enhancing pre-mRNA processing (Hur et al., 2020). Cyclin E/Cdk2-dependent phosphorylation of Mxc (ph-Mxc) can be detected in *Drosophila* HLBs using the MPM-2 monoclonal antibody (Calvi et al., 1998; White et al., 2007, 2011). ph-Mxc is not present in cells with inactive Cyclin E/Cdk2, and loss of Mxc phosphorylation at the end of S phase correlates with the down regulation of histone RNA. We hypothesized that Mute down regulates histone RNA by antagonizing the phosphorylation of Mxc. To test this hypothesis, we stained embryos with MPM-2 antibodies in combination with EdU labeling and RNA-FISH to the RD histone genes. In wild type embryos we observed concurrence among ph-Mxc, EdU, and both nascent RD histone RNA in the HLB and RD histone RNA in the cytoplasm (Fig. 6A, top, arrows). In *mute^1281^* embryos, we also observe cells that do not label with EdU but contain both nascent RD-histone RNA in the HLB and RD-histone RNA in the cytoplasm (Fig. 6A, bottom, double arrows). Strikingly these cells also contain ph-Mxc within the HLB, indicating that upon loss of Mute, Cyclin E/CDK2 phosphorylation of Mxc persists into G2, likely leading to the aberrant expression and accumulation of histone RNA in cells outside of S phase. These observations suggest a role for Mute as a regulator of Mxc phosphorylation which in turn controls full length histone expression and accumulation (Fig. 6C).

**Figure 6.**
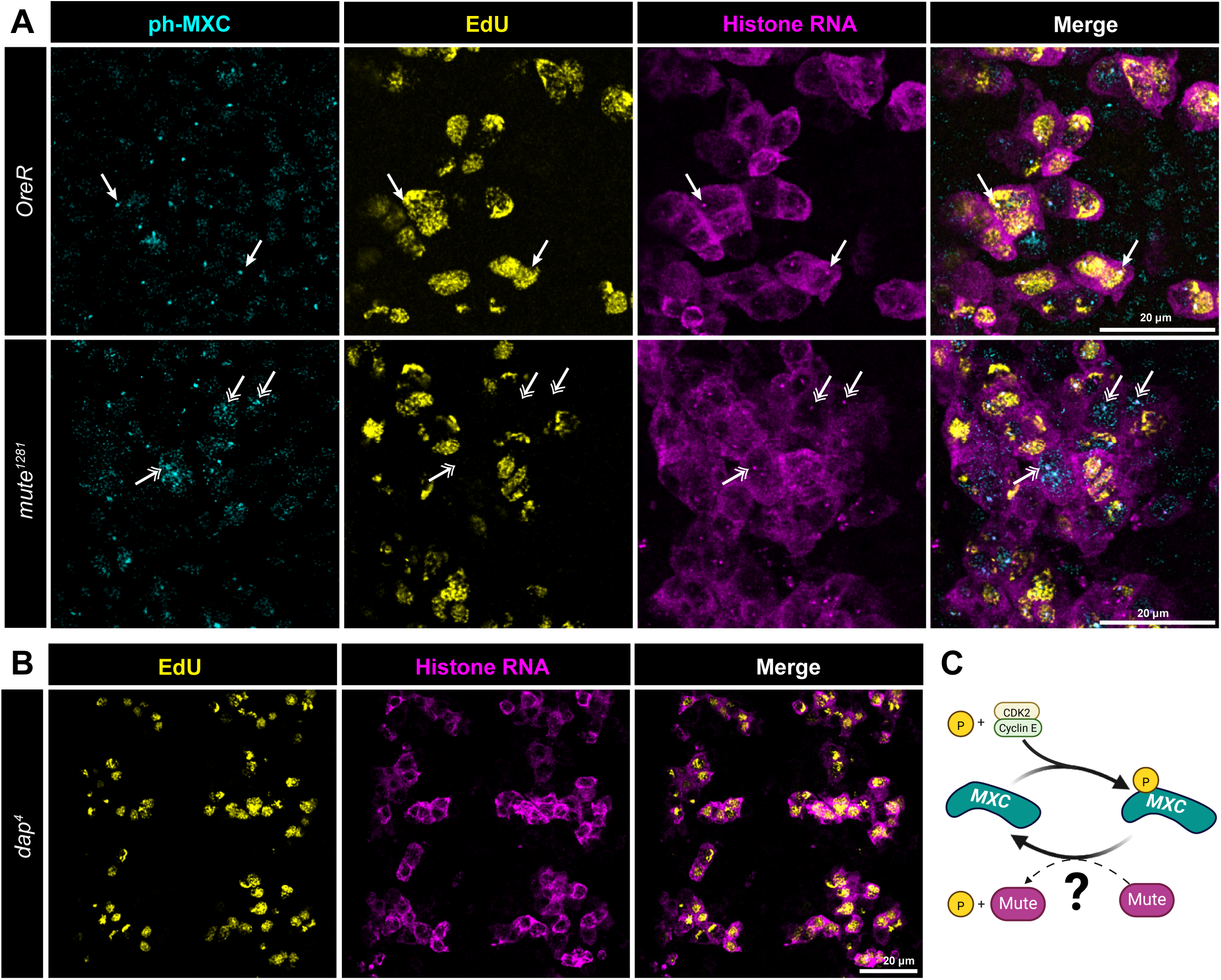
*mute* mutant cells fail to terminate Mxc phosphorylation **A)** VNC of *OregonR* and *mute^1281^* null embryos pulse labeled with EdU (yellow) and stained with MPM-2 antibodies recognizing phosphorylated Mxc (cyan) and hybridized with the CDS probe (magenta). Single and double arrows show cells with phosphorylated Mxc and nascent RD histone gene transcription in the HLB in EdU-positive and EdU-negative cells, respectively. Scale bar = 20μm. **B)** VNC of *Dap^4^* null mutant embryo pulse labeled with EdU (yellow) and hybridized with the CDS probe (magenta). Scale bar = 20μm. **C)** Proposed Model of Mute counter acting Cyclin E/Cdk2-dependent Mxc phosphorylation.

Mute has no catalytic domains and likely serves as an adaptor protein by binding directly to Mxc/NPAT and other components within the HLB (Bodner et al., 2026; Ingham et al., 2025; White et al., 2011; Yang et al., 2014). We considered the possibility that Mute might recruit an inhibitor of Cyclin E/Cdk2. *Drosophila* has a single p21-like Cyclin E/Cdk2 inhibitor encoded by the *dacapo* gene (Dap). Dap is first expressed zygotically in the embryo during interphase of cell cycle 16 to inactivate Cyclin E/CDk2 and trigger G1 arrest in cyclin 17 within the epidermis (de Nooij et al., 1996; Lane et al., 1996; Sauer et al., 1995). To determine whether Dacapo regulates RD histone RNA accumulation in the VNC we combined EdU labeling with RNA-FISH in *dap^4^* null mutant embryos (Fig. 6B). Loss of Dap results in inappropriate EdU labeling of stage 12 epidermal cells accompanied by ectopic expression of RD histone genes due to failure to down regulate Cyclin E/Cdk2 and enter G1 phase during cycle 17 (Lane et al., 1996). By contrast, in stage 14 *dap^4^* mutant embryos we observed no change in the pattern and concurrence between EdU labeling and histone RNA accumulation in the VNC (Fig. 6B). We conclude from these data that Mute recruitment of Dap to the HLB is not the primary mechanism for downregulating RD histone gene expression in the VNC.

### Mute is required for proliferation of wing imaginal disc cells

Disruption of both H1 and core RD histone gene expression is likely to affect cell proliferation. In addition, the Mute ortholog Gon4l is required for cell proliferation in zebrafish, mice, and human cancer cells (Agarwal et al., 2016; Barr et al., 2017; Budine et al., 2020; Lim et al., 2009). To determine whether loss of Mute function affects cell proliferation in *Drosophila*, we used the FLP/FRT system to generate clones of *mute^1281^* mutant cells in third instar wing imaginal discs, the precursors of the adult wing. We used heat-shock induced expression of the FLP recombinase to trigger mitotic recombination of chromosome 2R at the FRT42D site, which is centromere proximal to the *mute* locus (Fig. 7A). The chromosome carrying wildtype *mute* is marked by a distal insertion of Ubiquitin-GFP. Consequently, stochastic occurrence of mitotic recombination following heat shock induced FLP expression results in two daughter cells of differing genotype: one homozygous wildtype for *mute*, marked by 2x expression of GFP, and one homozygous for the *mute^1281^* null mutation, marked by no GFP. Equal proliferation of these two daughter cells will result in “twin spots” of 2x GFP wild type cell clones adjacent to GFP negative *mute* null cell clones in a background of 1x GFP heterozygous cells (Fig. 7A,C, D).

**Figure 7.**
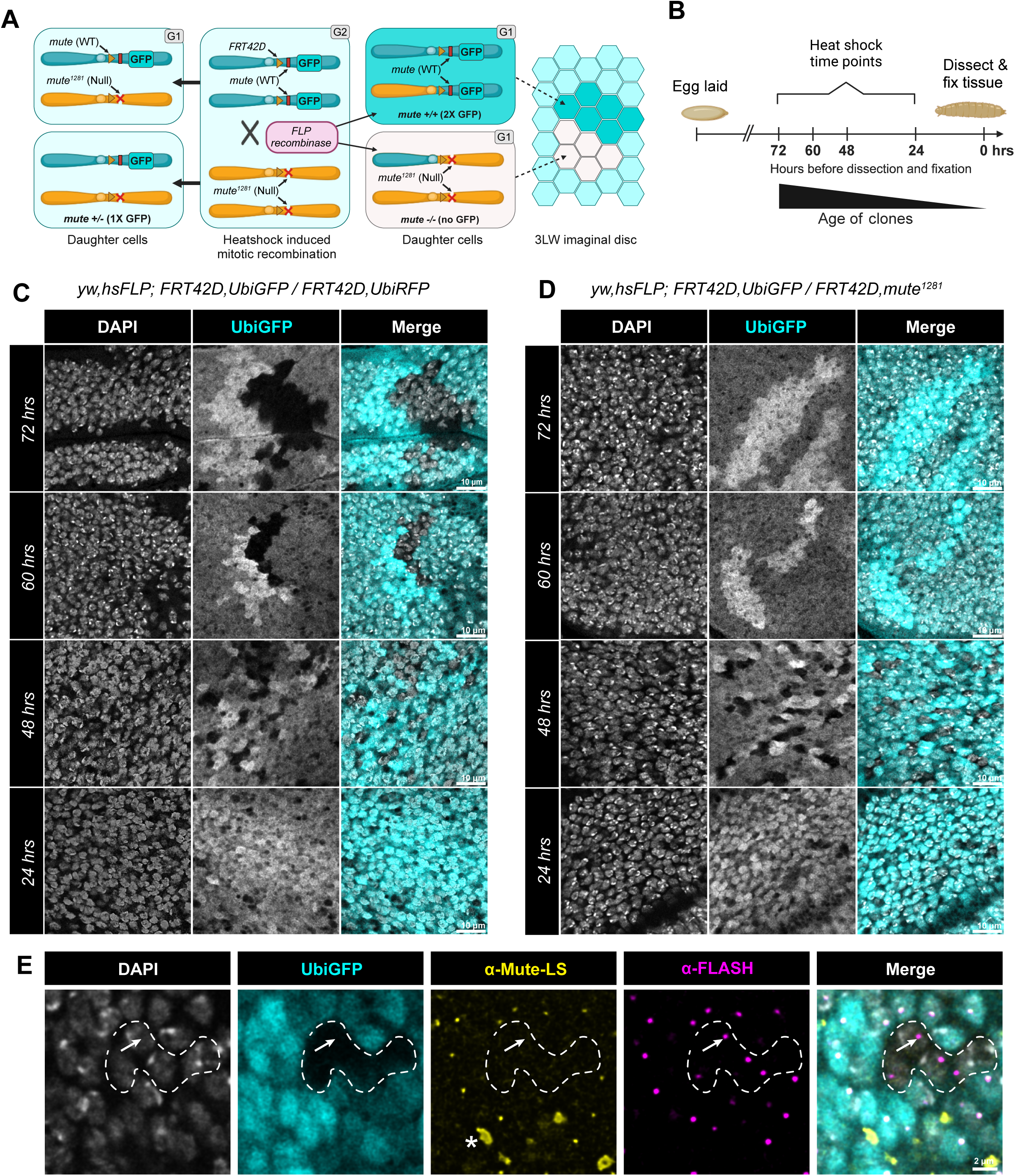
*mute* mutant wing imaginal discs cells do not proliferate **A)** Schematic of the mitotic recombination method used to make twin spot clones of *mute^1281^* null mutant cells adjacent to *mute* wild type cells. **B)** Schematic of larval development and when mitotic recombination was induced. **C)** Twin spots resulting from mitotic recombination induced at the indicated times before dissection using a control wild type chromosome. Scale bar = 10μm. **D)** Twin spots resulting from mitotic recombination induced at the indicated times before dissection using a *mute^1281^* null mutant chromosome. Note that by 60 hours before dissection no *mute* mutant clones are detectable. Scale bar = 10μm. **E)** *mute^1281^* null mutant cell clone (dotted line) induced by mitotic recombination and stained with antibodies recognizing Mute (yellow) or FLASH (magenta). Note the absence of Mute accumulation in the HLB in the *mute^1281^* mutant cells. Asterisk denotes unknown extracellular aggregates in Mute channel, commonly seen when making mitotic clones of *mute^1281^* cells. Scale bar = 2μm.

The wing disc forms during embryogenesis from a group of around 30 cells and then grows during larval development to about 35,000 cells at the wandering third instar larval stage (Tripathi & Irvine, 2022). We induced mitotic recombination at different times during larval development by heat-shocking at 72, 60, 48, and 24 hours prior to dissecting wing discs at the wandering third instar larval stage (Fig. 7B). Using this strategy, we can get a range of different sized clones and determine if the *mute* mutant cells grow at a comparable rate to wildtype. When this protocol was applied to *FRT42D, Ubi-GFP/FRT42D, Ubi-RFP* homozygous wild type control animals, we identified obvious twin spots with equally sized GFP^+^ and GFP-negative clones at 72, 60, and 48 hours after heat shock with the 72 hour time point resulting in the largest clones (Fig. 7C, top). Dissection 24 hours after heat shock resulted in very small wild type clones that were barely visible (Fig. 7C, bottom). Wing discs from *FRT42D, Ubi-GFP/FRT42D, mute^1281^* third instar larvae contained GFP-negative clones homozygous for *mute^1281^* in the 48 hours before dissection samples (Fig. 7D, 3^rd^ row), but not in the 60 or 72 hours before dissection samples which only contained clones of wild type GFP^+^ cells (Fig. 7D, 2^nd^ and 1^st^ rows, respectively). These data indicate that *mute* null cells have a proliferation defect and are ultimately eliminated during growth of the tissue, a well described phenomenon in wing discs (Martín et al., 2009; Morata & Ripoll, 1975; Simpson, 1979). At 48 hours after heat shock no homozygous *mute^1281^* clones larger than 8 cells were observed, and these cells did not contain detectable levels of Mute in the HLB, marked by FLASH (Fig. 7E, arrow). As in the embryonic VNC, this result indicates that Mute is not required for HLB formation.

To determine if *mute^1281^* cells are eliminated from the wing disc due to competition with neighboring wildtype cells, we used the heat shock FLP/FRT system to generate *mute^1281^* mitotic clones in a *M(2)53(1) Minute* mutant background (Fig. S5A), which is slower growing and will reduce competition (Fig. S5B)(Bridges & Morgan, 1923; Lambertsson, 1998; Morata & Ripoll, 1975). If *Mute^1281^* cells proliferate slowly rather than becoming arrested, then we expect to find larger *Mute^1281^* clones in the Minute background at 60 and 72 hours after heat shock compared to a wildtype background as in Fig. 7. Homozygous wild type *mute* control clones proliferated well in the heterozygous *Minute* background with the largest clones appearing at 72 hours after heat shock induction of mitotic recombination (Fig S5C). As expected, clones homozygous for the *Minute* mutation did not grow past single cells. Although *mute^1281^* clones (marked by the absence of RFP) in the *Minute* background persisted at 48 and 60 hours, they did not grow larger than 8 cells (Fig. S5D) as we observed in the wild type background. The inability of the *mute^1281^* clones to grow large even in a reduced competition background shows that *mute^1281^* cells have a proliferation defect rather than simply being slow growing. The upper limit on the size of the *mute^1281^* clones suggests that wing disc cells lacking Mute arrest after a few cell divisions, differing from the cells of the embryonic VNC.

## Discussion

### Mute plays dual roles in *Drosophila* RD histone gene regulation

In this study, we show that Mute functions to restrict the expression of all RD-histone genes to S phase, while simultaneously having differential effects on the maximum transcriptional output during S phase of *H3* and *H4* versus *H1*, *H2a*, and *H2b*. Upon loss of Mute, neural progenitor cells within the embryonic VNC inappropriately transcribe and accumulate RD histone mRNA in G2 phase. This phenotype is accompanied by the occurrence of phosphorylated Mxc, suggesting that Mute either mediates the dephosphorylation of Mxc or the inhibition of Cyclin E/Cdk2 activity within the HLB at the end of S phase. We favor the former, as Mute antagonizes phosphorylation of Mxc independently of the Cyclin E/Cdk2 inhibitor Dacapo. Cyclin E/CDK2 controls transcription of *Drosophila* RD histone genes by stimulating release of promoter-proximal pausing of RNA pol II (Kemp et al., 2025), suggesting that Mute might act as a negative regulator of transcriptional pause release.

Mute is a large protein with multiple intrinsically disordered regions (IDRs) and two centrally located PAH domains and a C-terminal SANT domain. The PAH domains share both sequence and predicted structural homology with Sin3a, a well-studied protein that serves as a hub for the assembly of transcriptional repressor complexes (Bugge et al., 2021; Sahu et al., 2008). Gon4l, the vertebrate ortholog of Mute, has been found in a complex with Sin3a, YY1, and HDAC1 that can repress expression of a reporter construct in mouse cells (Lu et al., 2011). Very few Mute binding partners have been reported. One is the HLB scaffold Mxc, which can bind Mute’s C-terminal SANT domain *in vitro*, an interaction conserved in the human orthologs Gon4L (aka YARP) and NPAT (Yang et al., 2014). Whether the Mute-Mxc interaction is important for control of RD histone gene expression has not been investigated. Mute co-immunoprecipitates with the *Drosophila* protein Winged-Eye (WGE) from cultured S2 cell extracts (Ozawa et al., 2016). Like Mute, WGE localizes to the HLB and steady state amounts of core histone mRNA are elevated in *wge* mutants (Ozawa et al., 2016). Thus, WGE may partner with Mute to regulate core histone genes. Identification of additional Mute binding partners along with analysis of Mute’s various functional domains will be crucial in elucidating its molecular mechanism of action.

In addition to WGE, other proteins in *Drosophila* and other organisms have been reported to act as negative regulators of histone gene expression, including Abo (now Ao), HIRA, HERS, WEE1, ATR, and histone H4 (Ahmad et al., 2026; Berloco et al., 2001; Ito et al., 2012; Mahajan et al., 2012; Marmolejo et al., 2026; Nelson et al., 2002; Ozawa et al., 2016; Roth et al., 2025). Of these, Mute, WGE, ATR, and H4 have cytological evidence of enrichment in HLBs. A recent study demonstrated that Ao does not localize to the HLB or control steady state levels of RD histone mRNA (Takenaka et al., 2026). Phosphorylation of Tyr37 of H2B by WEE1 at histone genes in mouse cells precludes binding of NPAT and RNA pol II, and inhibition of WEE1 results in failure to down regulate histone expression at the end of S phase (Mahajan et al., 2012). Mammalian HIRA has long been posited as a negative regulator of histone transcription, similar to the function of its *S. cerevisiae* homologues Hir1p and Hir2p (Hall et al., 2001; Nelson et al., 2002). In mammalian cells, ectopic expression of HIRA represses histone gene expression and blocks DNA synthesis (Nelson et al., 2002), but it’s well-documented function as a chaperone for replication-independent histone H3.3 (Choi et al., 2024; Ricketts & Marmorstein, 2017) suggests that any effect of HIRA on histone gene expression may be indirect. Finally, two recent studies provide evidence that Histone H4 accumulation in the HLB provides negative feedback regulation to RD histone gene expression in both flies and human cells (Ahmad et al., 2026; Roth et al., 2025). Some of these factors may work in concert with Mute/Gon4L to coordinate cell cycle inputs and repress RD histone transcription at the end of S phase.

In marked contrast to its role in repressing core RD histone gene expression outside of S phase, we found that Mute is also required for full expression of H1, H2a, and H2b mRNA, a result analogous to recent observations made in vertebrates regarding H1. Like Mute, zebrafish Gon4l is required for H1 expression (Williams et al., 2018). The PAH domains of human Gon4L bind to CRAMP1, which accumulates in HLBs and localizes to the promoters of transcriptionally active H1 genes. CRAMP1 activates H1 gene expression but does not regulate core histone genes (Bodner et al., 2025; Ingham et al., 2025; Matthews et al., 2025). The *Drosophila* ortholog of CRAMP1 is encoded by the *cramped* gene, which is also required for expression of H1 but not core RD histone gene expression (Gibert & Karch, 2011). Whether human Gon4l negatively regulates core RD histone genes has not been addressed. Together with our work, these recent findings on CRAMP1 suggest a conserved role for Mute/Gon4L in regulating *H1* expression mediated by an interaction with Cramped/CRAMP1. There is precedence for independent control of *H1* and core RD histone genes in *Drosophila*, as the TATA binding protein TRF2 is required for *H1* expression, but not core RD histone genes (Guglielmi et al., 2013; Isogai et al., 2007).

These observations raise an interesting question of how Mute can uniformly affect RD histone genes in one manner but have a differential effect in another. One possibility is the existence of multiple Mute complexes within the HLB that have specificity for either the *H3-H4* bidirectional promoters or the *H2a-H2b* bidirectional promoters and the *H1* promoter within the *Drosophila* RD histone gene unit. Another possibility is that one Mute complex affects all RD-histone genes in the same manner (e.g. activation), but the persistence of histone gene transcription into G2 leads to an independent upregulation of H4 and H3 in a feedback mechanism to repress aberrant expression. Our work highlights the importance of examining multiple facets of RD histone gene expression when assessing purported regulators including: (1) whether maximal, per cell transcriptional output is altered; (2) whether aberrant expression occurs outside of S-phase; (3) whether bulk histone mRNA levels are changed within a given tissue or organism; and (4) whether all replication-dependent histone genes are uniformly affected.

### Loss of Mute or Gon4l leads to pleiotropic effects on cellular proliferation and development

Numerous studies found that Mute and Gon4l participate in a variety of developmental and pathological processes. Mute stands for *muscle wasted* and was first identified in *Drosophila* as an embryonic lethal mutation causing defects in muscle differentiation and loss of muscle mass (Bulchand et al., 2010). Similarly, an unbiased forward genetic screen found *mute* is required for development of Ap4/FMRFa neurons in the embryonic VNC (Bivik et al., 2015). Mutation of vertebrate Gon4l disrupts immune cell development in mice (Lu et al., 2010), neurogenesis in cultured rat cells (Li et al., 2024), notochord formation and cardiomyocyte maintenance in zebrafish (Budine et al., 2020; Williams et al., 2018), and causes dwarfism in cattle (Schwarzenbacher et al., 2016). A recent human clinical study found two separate cases of bi-allelic *Gon4l* variants leading to microcephaly, craniofacial abnormalities, and developmental delay (Li et al., 2024). Mutations in other HLB proteins such as NPAT have been linked to several diseases including cancer (reviewed in (Geisler et al., 2023)), and elevated RNA polymerase II at histone genes correlates with poor clinical outcomes of numerous cancers (Henikoff et al., 2025).

What could be a common explanation for all these observations? One possibility is that Gon4l/Mute have a variety of gene targets that result in pleiotropic effects on developmental processes when absent. There is support for this model from studies in zebrafish and mice (Budine et al., 2020; Tsai et al., 2020). In contrast, our cell biological and genomic data indicate that Mute, like Mxc, is present solely in the HLB and localizes only to RD histone genes. This pattern of enrichment on the genome is similar to other ChIP-seq and CUT&TAG datasets of Mxc and Mute (Ahmad et al., 2026; Hodkinson et al., 2024). So how can proteins that only regulate RD histone genes have such pleiotropic effects on development when mutated? A common thread may be disruption of cell proliferation, as we observed in *mute* mutant wing imaginal discs. Mutation of Gon4l prevents B cell proliferation in mice (Barr et al., 2017) and disrupts proliferation of human cancer cells (Agarwal et al., 2016). Mxc mutations disrupt normal cell division in several *Drosophila* tissues (Kurihara et al., 2020; Sang et al., 2022; Tanabe et al., 2019).

However, another result of mis-regulating RD histone gene expression could be disruption of chromatin causing widespread changes in gene expression, the latter of which we observed in *mute^1281^* embryos by RNA-seq. A good example of this possibility is Polycomb-mediated gene repression. Mxc stands for *multi sex combs* and was originally identified as a classic *polycomb*-like mutant (Santamaria & Randsholt, 1995). Mxc and Cramped are the only Polycomb group (PcG) genes that genetically interact with other PcG genes without ever being shown to physically interact with any PcG proteins (Gibert & Karch, 2011; Kassis et al., 2017; Santamaria & Randsholt, 1995). These observations suggest that normal histone gene expression is needed to establish a repressed chromatin configuration at PcG-regulated genes. Indeed, CRAMP1-directed H1 expression is necessary for PRC2-based repression in human cells (Matthews et al., 2025), and reduction of RD histone gene dose in *Drosophila* disrupts Pc function (McPherson et al., 2023). This phenomenon may extend to constitutive heterochromatin, as reduced histone gene copy number suppresses position effect variegation (Moore et al., 1983). Thus, tight control of histone gene expression is needed for chromatin configurations that support normal gene expression, and consequently the proper differentiation and function of many cell types.

## Acknowledgements

We thank Cameron Prince for help generating the mCherry2-Mute line, the UNC High Throughput Sequencing Facility for sequencing CUT&RUN and RNA-Seq samples, the UNC Proteomics core for producing protein A/G-MNase for CUT&RUN, Markus Nevil for creation of the custom dm6-chrHis genome, and the J. Skeath (WashU), C. Doe (Oregon), and Y.N. Jan (UCSF) labs for sharing guinea pig anti-Deadpan and other antibodies to neural markers. This work was supported by NIH F31HD113267 to M.S.G. and NIH R35GM145258 to R.J.D.

## Methods and Materials

### Fly strains and husbandry

The following *D. melanogaster* strains were used in this study: *OregonR*. *yw; mute^1281^/ cyo-twi-gfp* (Bulchand et al., 2010) (BDSC: 34055). *yw;;nos-Cas9(III-attP2)* (BestGene, Inc.). *yw; mCherry2-Mute* (this study). yw; Mute-DF/cyo-twi-GFP (w[1118]; Df(2R)Exel6065, P{w[+mC]=XP-U}Exel6065/CyO; BDSC: 747). *w^1118^; mute^1281^/ cyo-twi-gfp; Ub-FUCCI(III) (ub-GFP-E2F1 ^#5^ ub-mRFP1-NLS-CycB ^#12^* from (Zielke et al., 2014) BDSC: 55124). *yw;MiMIC-Dpn(GFP)/Sm6a* (BDSC: 59755). *w[*]; dap[4]/CyO, P{ry[+t7.2]=ftz-lacB}E3* (BDSC: 6639). *yw,hsFLP;FRT42D, UbiGFP/Cyo* (from BDSC: 5626). *yw,hsFLP;FRT42D, UbiRFP* (from BDSC: 35496). *yw,hsFLP;FRT42D,Mute^1281^/Cyo-twi-GFP*. *yw,hsFLP;FRT42D, UbiRFP, M(2)53(1)/Cyo* (from BDSC: 5698). Embryos were genotyped during imaging using fluorescent cyo-twi-gfp balancers stained for α-GFP.

### Antibodies

Secondary antibodies used in this study include goat anti-rabbit Atto 488 (Sigma Cat #18772), goat anti-guinea pig Alexa 488 (Invitrogen), goat anti-Mouse IgG1 Alexa Fluor 488 (Invitrogen A21121), goat anti-Rabbit IgG Alexa+ Fluor 555 (Invitrogen A32732), goat anti-guinea pig Alexa Fluor 555 (Invitrogen A21435), goat anti-Mouse IgG1 Alexa Fluor 555 (Invitrogen A-21127), goat anti-guinea pig Alexa Fluor 647 (Invitrogen A21109), or goat anti-rabbit Alexa fluor 680 (Invitrogen A21450) each diluted at 1:1000 in 0.1% PBST. Primary antibodies and their specific dilutions are listed below:

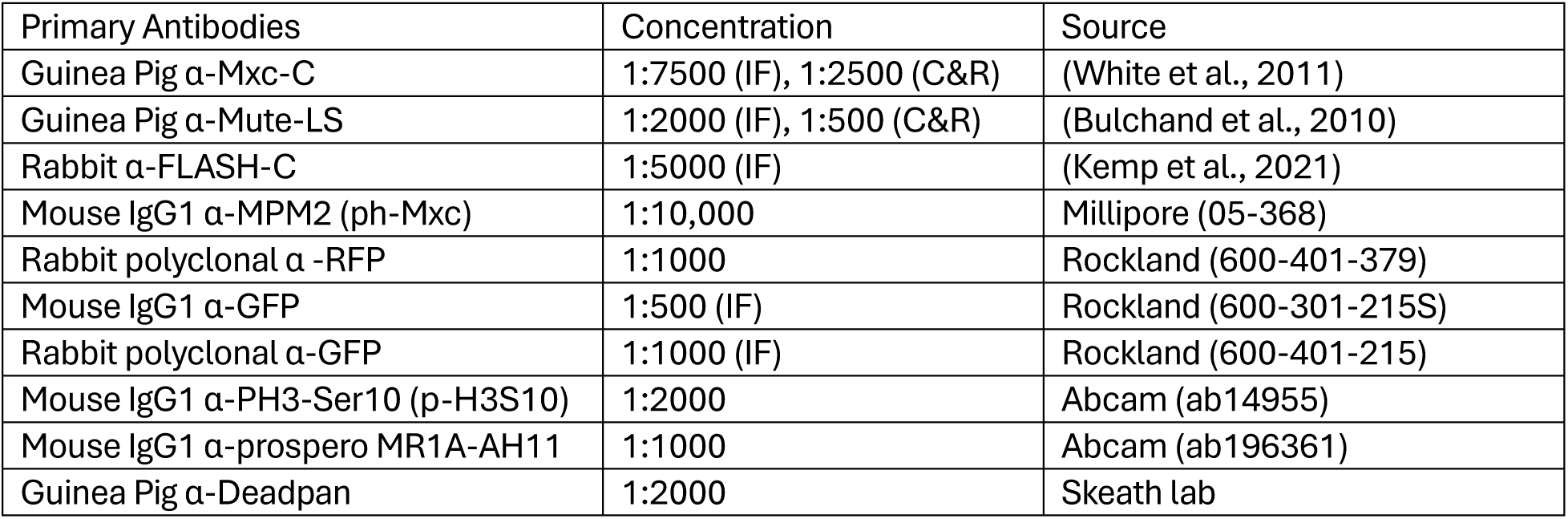

| Primary Antibodies | Concentration | Source |
| --- | --- | --- |
| Guinea Pig α-Mxc-C | 1:7500 (IF), 1:2500 (C&R) | (White et al., 2011) |
| Guinea Pig α-Mute-LS | 1:2000 (IF), 1:500 (C&R) | (Bulchand et al., 2010) |
| Rabbit α-FLASH-C | 1:5000 (IF) | (Kemp et al., 2021) |
| Mouse IgG1 α-MPM2 (ph-Mxc) | 1:10,000 | Millipore (05-368) |
| Rabbit polyclonal α -RFP | 1:1000 | Rockland (600-401-379) |
| Mouse IgG1 α-GFP | 1:500 (IF) | Rockland (600-301-215S) |
| Rabbit polyclonal α-GFP | 1:1000 (IF) | Rockland (600-401-215) |
| Mouse IgG1 α-PH3-Ser10 (p-H3S10) | 1:2000 | Abcam (ab14955) |
| Mouse IgG1 α-prospero MR1A-AH11 | 1:1000 | Abcam (ab196361) |
| Guinea Pig α-Deadpan | 1:2000 | Skeath lab |

### Generation of mCh2-Mute CRISPR fly line

To generate the endogenously tagged Mute fly line yw;;nos-Cas9(III-attP2) embryos were injected (BestGene, Inc.) with two plasmids, one containing a guide sequence targeting the 5’ end of the long isoform of the Mute gene and one containing a repair template. The guide sequence was designed using the flycrispr target finder (http://targetfinder.flycrispr.neuro.brown.edu/) and the Drosophila melanogaster reference genome, r_6. The Mute target sequence was GTCCGACTTCTTTGCACTGCT_CGG. The guide plasmid was created via PCR using the following primers to pU6-2-BbsI-gRNA: Rev primer GAAGTATTGAGGAAAACATACCTATATAAATG, and FWD primer GTCCGACTTCTTTGCACTGCTGTTTTAGAGCTAGAAATAGCAAG. The PCR was carried out as described in <u>U6-gRNA (chiRNA) cloning – flyCRISPR</u>. The repair template was synthesized by Genewiz and was designed with a *D. melanogaster*codon optimized version of mCherry2 that was inserted after the second codon of the Mute gene separating the PAM sequence from the target sequence of the guide. The full sequence of the plasmid is as follows.

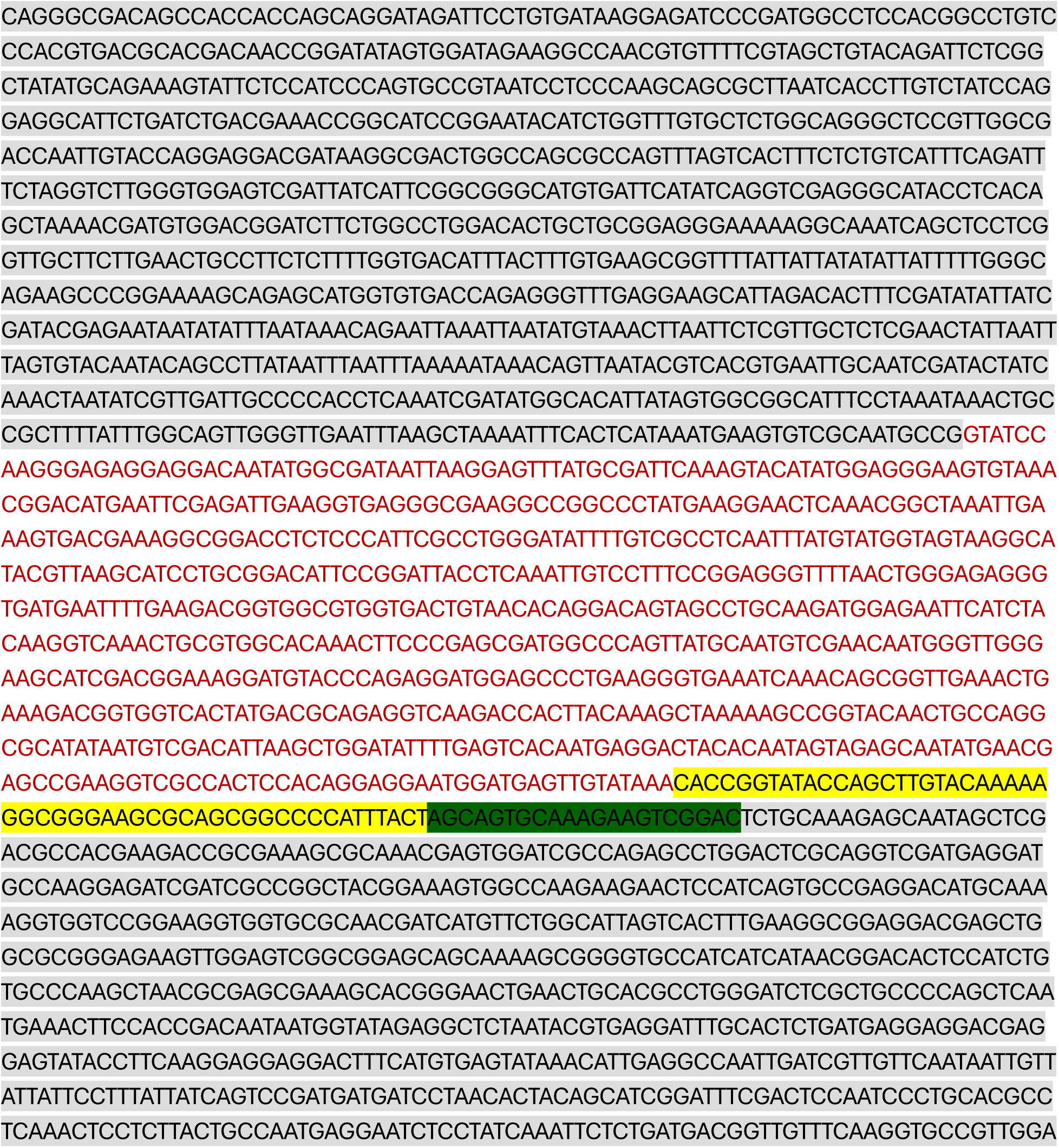
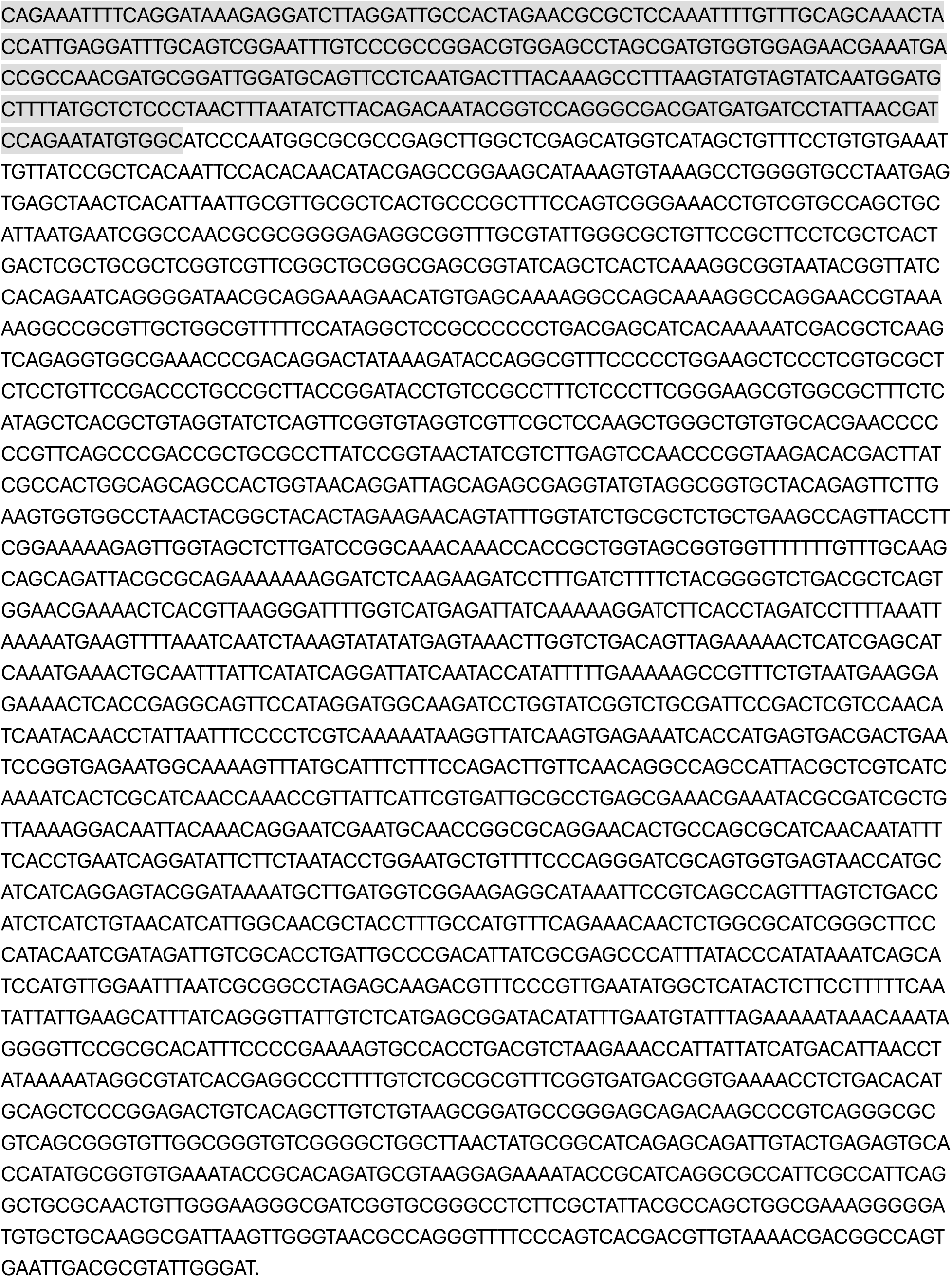

The Homology arms are highlighted in grey, the mCherry2 sequence is in red, the linker used is highlighted in yellow, and the guide target sequence is highlighted in green. Adult flies from injected embryos were crossed to yw;Pin/Cyo as single crosses and 10 Cyo progeny were backcrossed to yw; Pin/Cyo as single crosses again. These flies were screened for insertion via PCR with the following primers.

FWD_TTGCCCCACCTCAAATCG, and Rev_CGATCGATCTCCTTGGC. Cyo progeny from positive vials were self-crossed and homozygous mCherry2-Mute lines were established.

### Generation of RNA-FISH probes

Custom Stellaris RNA-FISH probes (Orjalo & Johansson, 2016; Raj et al., 2008) were generated using Stellaris® RNA-FISH Probe Designer (LGC Biosearch Technologies, Petaluma, CA). The CDS probe set, to the H2a, H2b, H3 and H4 coding region and H3 only CDS were described previously (Hur et al., 2020; Kemp et al., 2025). The H1 probe set, which covers only the 3’ portion of H1, starting 200bp after the TSS and the H2a probe set which covers the 5’UTR, entire CDS, and 3’UTR, were designed for this study and the oligonucleotides and sequences used to design them are listed below.

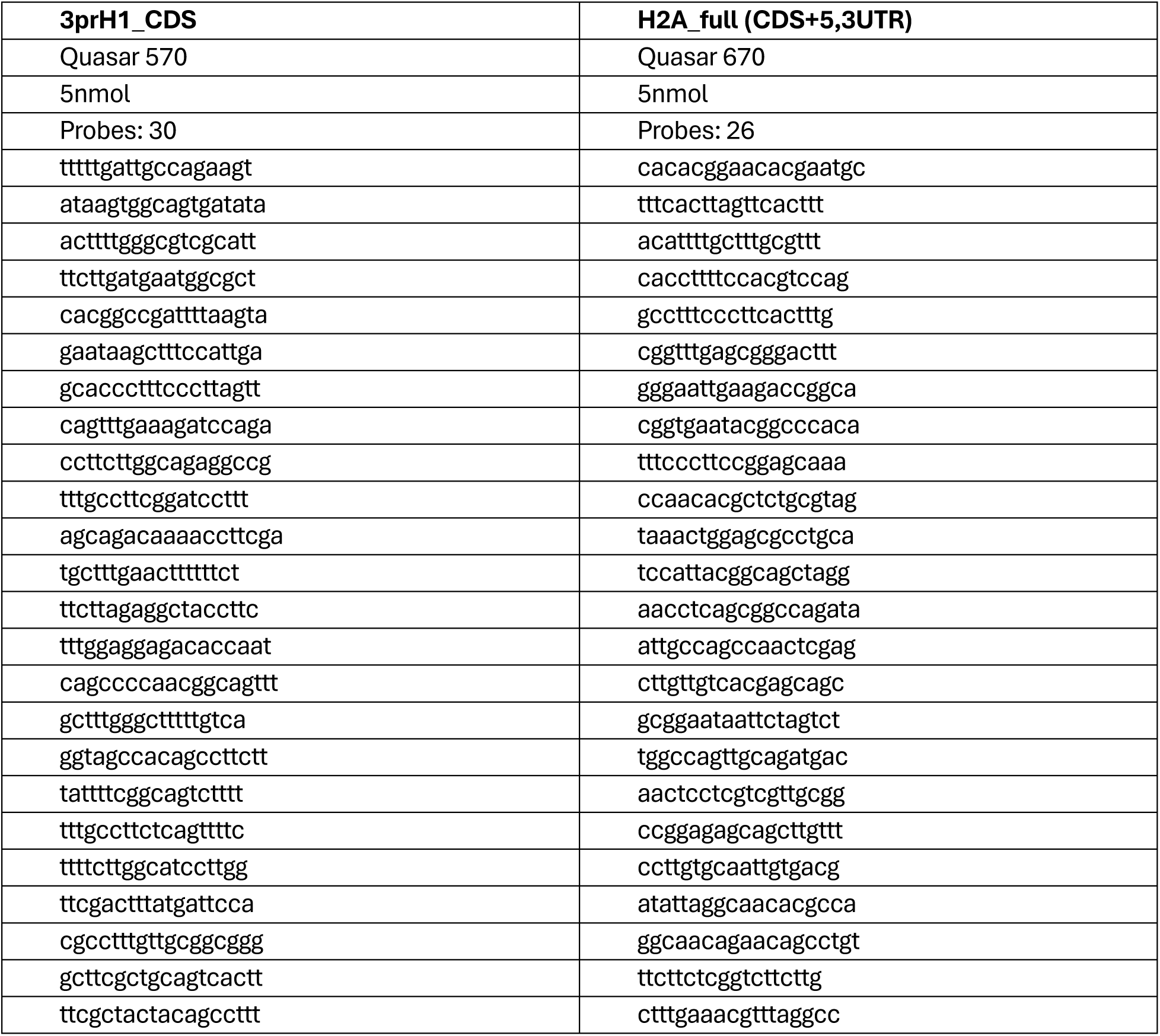
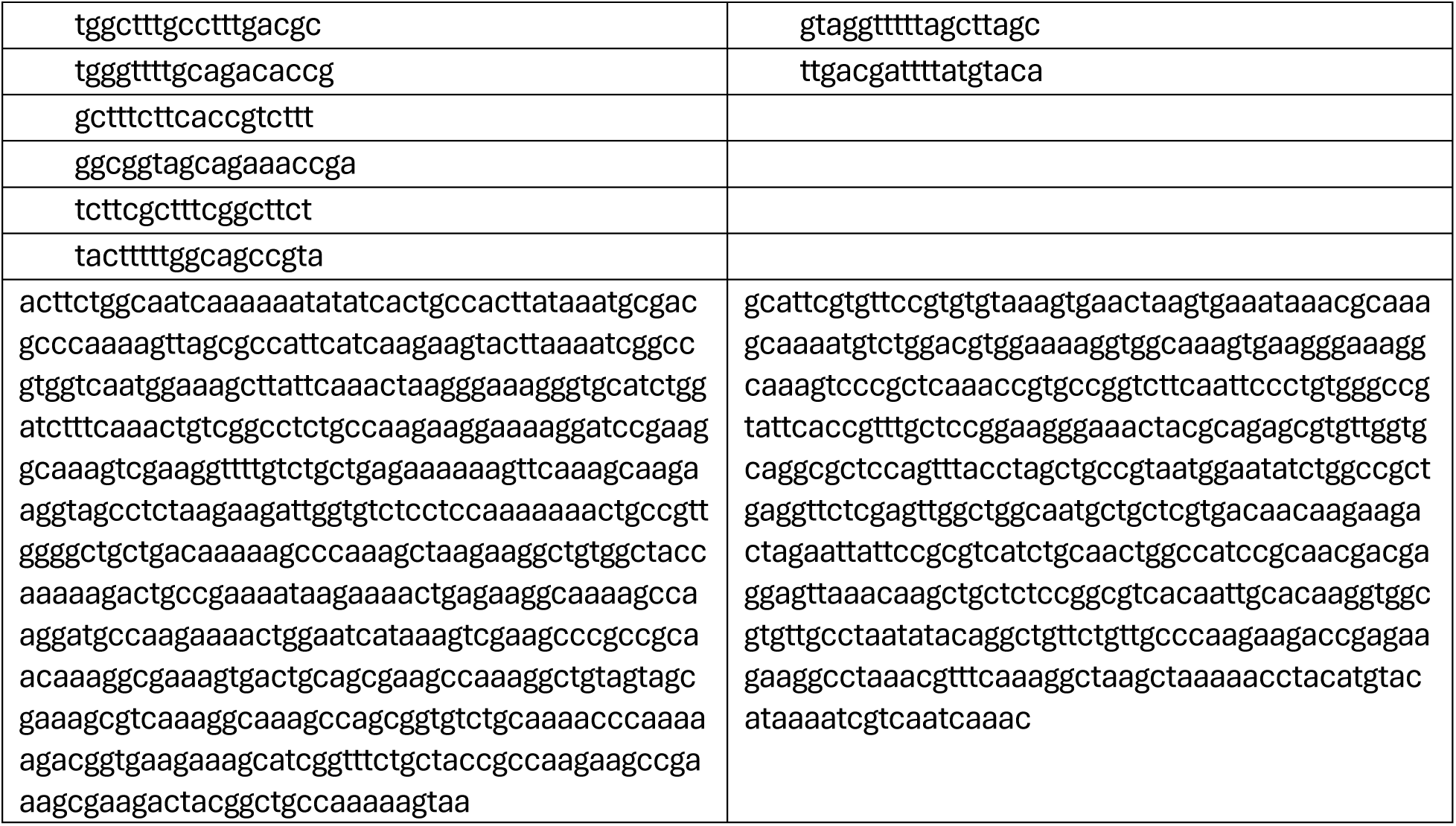

### Immunofluorescence and RNA-FISH of embryos

Embryos 9-11hrs (Stage 14 were collected on apple juice agar plates at 25°C and aged in the dark at 25°C. Embryos were removed from the plate into a 1.7mL Eppendorf tube and washed once using Embryo Wash buffer (120mM NaCl, 0.07% Triton-X100). Embryos were dechorionated using 50% freshly diluted bleach for 5 minutes and then washed twice with embryo wash buffer. Embryos were fixed in 0.5mL 4% paraformaldehyde and 0.5 mL heptane for 20 minutes while nutating at room temp. The aqueous phase (paraformaldehyde) was removed and 0.5mL Methanol was added. The tube was shaken vigorously for 30 seconds to remove the embryos’ vitellin membranes. Methanol and heptane were immediately removed and replaced with 0.5mL fresh methanol. Embryos were either stored at −20°C or rehydrated in PBST immediately. To rehydrate, embryos were nutated in equal parts methanol: 0.1% PBST, ex1:2 ratio of methanol: 0.1% PBST, and finally 1.0mL 0.1% PBST, each for 10 minutes at room temp. Embryos were then stained or stained and hybridized with RNA-FISH probes as previously described (Kemp et al., 2025) and imaged on a Leica SP8 confocal microscope with LIGHTNING system.

### 5-ethynyl-2’-deoxyuridine (EdU) labeling, Immunofluorescence, and RNA-FISH in embryos

Aged embryos were collected into a nylon mesh basket using embryo wash buffer and washed once more. The mesh basket was submerged in 50% bleach for 4 minutes to dechorionate embryos and then rinsed for 2 minutes under running tap water to remove bleach. The mesh basket was then blotted dry and allowed to air dry for 3-5 minutes depending on the humidity. A dissecting scope was used to monitor drying of the embryos. Once dry, the basket was submerged in octane for no more than 3 minutes to permeabilize the vitellin membrane. The embryos were spread into a monolayer by gently bouncing the basket. After 3 minutes, the basket was removed and blotted from the bottom. A dissecting scope was then used to watch the octane evaporate off the embryos. Just as almost all octane was evaporated, 4Ml of 0.25mg/mL EdU in Grace’s media was gently poured on top of the embryos, without dislodging the monolayer of embryos from the mesh. Embryos were incubated in EdU for 30 minutes. One minute prior to the end of the incubation, the basket was removed into a watchglass of heptane to allow transfer of the embryos into a 1.7mL Eppendorf tube. To fix, 0.5mL of 4% paraformaldehyde was added to the embryos already in a 0.5mL volume of heptane and then nutated for 20 minutes at room temp. Embryos were then de-vitellinized and rehydrated as described above and stained and hybridized as described previously (Kemp et al., 2025). Slides were imaged on a Leica SP8 confocal microscope with LIGHTNING system.

### Image Analysis and Quantification

#### Image Analysis

All image analysis was performed using FIJI software (Schindelin et al., 2012).

#### Quantification of cell classes

To quantify the co-occurrence of EdU and Histone mRNA in the VNC of stage 14 embryos, 1.8 µm deep max intensity projections were labeled manually using the FIJI Multipointer tool with DAPI as a guide. Edu positive nuclei, which always co-occurred with RNA-FISH, were labeled, counted, and stored as a selection which could be overlayed on the image. This overlay was then used to label, count, and store the nuclei which contained RNA-FISH signal but no EdU. This selection was pseudocolored and overlayed with the EdU selection to make sure no nuclei were counted twice. This was performed on 4 embryos for each genotype and plotted using GraphPad Prism. The same method was used to classify and count Cyclin-B and phos-H3 Serine 10 positive cells and their overlap with histone RNA as well as the overlap of deadpan, prospero, and histone RNA positive cells in the VNCs of *mute^1281^* homozygous and heterozygous siblings or OreR and *mute^1281^* embryos respectively (n=4 for all but p-H3S10 where n = 3 embryos per genotype). The labels and counts were saved as selections and plotted using GraphPad Prism.

#### Quantification of RNA-FISH signal

Pixel intensities of 1.8 µm deep z-stacks centered on the VNC were summed and then binned to generate histograms of pixel intensities across 100 µm^2^ images in FIJI. 5 biological replicates of each probe sets targeting histone genes H3, H2a, H1, and CDS were analyzed in each genotype. H3 and H2a were acquired from the same embryos (different channels), as were H1 and CDS, though this pairing was not relevant to the analysis. Histograms were collected with a bin size of 1 intensity unit. H3 and H2a histograms spanned bins 0–699, H1 and CDS histograms spanned bins 0–1399. All bins were retained for analysis. Between-genotype comparisons were performed for each of the four probe sets (OreR vs. mute1281 for H3, H2a, H1, and CDS) as unpaired analyses. Data was analyzed and plotted in R (v4.5.0).

### Mute and Mxc Cut&Run library preparation

Cut&Run in *OregonR* Wing imaginal discs was performed as previously described in (Stutzman et al., 2024). Three replicates per antibody were performed with 20 wing discs per sample. The following antibodies and concentrations were used: Guinea Pig Anti-Mxc-C used at 1:2,500, Guinea Pig Anti-Mute-LS used at 1:500. No secondaries were used. Pooled libraries were sequenced PE75 on a NextSeq2000 P1 flow cell.

### CUT&RUN Data Processing

Three replicates for each antibody and an IgG control were processed using the nf-core/cutandrun pipeline (v3.2.2; doi:10.5281/zenodo.5653535) (Ewels et al., 2020) executed via Nextflow (v24.04.2) using the Apptainer/Singularity container profile (unc_longleaf; https://github.com/nf-core/configs/blob/master/conf/unc_longleaf.config). The reference genome used was a custom dm6 genome (dm6-chrHis) where the partially assembled RD histone locus (*HisC*; chr2L:21,400,839-21,573,418) was removed and replaced with a single copy of the 5080bp histone gene repeat as its own chromosome (chrHis). Raw FASTQ files underwent adapter trimming and quality filtering using Trim Galore! (v0.6.7). Read quality was assessed by FastQC before and after trimming, and summary reports were aggregated using MultiQC. Trimmed reads were aligned to the custom genome using Bowtie2 (v2.5.4) with the following flags: --end-to-end --very-sensitive --no-mixed --no-discordant --phred33 -I 10 -X 700. Duplicate-marked, sorted BAM files from the nf-core pipeline were merged across biological replicates for each experimental condition using SAMtools (v1.23) Genome-wide signal tracks were generated from merged BAM files using deepTools (v3.5.6) bamCoverage to generate CPM normalization BigWigs (bin size = 2).

### Per-Histone-Gene-Copy (HCG) Normalization of chrHis Signal

The chrHis chromosome represents a single collapsed consensus sequence of the Drosophila histone gene array, which in the fly genome is present as a tandem repeat of approximately 100 copies. To correct for this copy-number artifact and produce signal values that are directly comparable in scale to single-copy genomic regions, all CPM signal across the entire chrHis chromosome was divided by 100, yielding per-histone-gene-copy (HCG) normalized signal using pyBigWig library in Python (v3.12.4). These HCG-normalized bigWig files were visualized in IGV (v2.19.04) and used for all subsequent quantitative deepTools analyses.

### CUT&RUN Analysis

Signal matrices were computed from the HCG-normalized bigWig files across all genes using deepTools (v3.5.6) computeMatrix. Gene bodies were scaled to 2000bp and included 500bp before the TSS and after the TES. The dm6 blacklist (v2) was applied to exclude signal from problematic regions (Amemiya et al., 2019). Genes were sorted in descending order of total signal, then clustered (--hclust 3) and plotted using deepTools plotHeatmap. Mean genome-wide signal was also computed in non-overlapping 5,080bp (chrHis size) bins across the genome using deepTools multiBigwigSummary. The dm6 blacklist (v2) was again applied to exclude signal from problematic regions. Manhattan plots of signal across the genome were generated in Python using matplotlib. Signal matrices of Mute signal across the five RD-histone genes as well as 450 up and 351 down regulated DEGs were computed using computeMatrix with the three sets of genes provided as BED files and again using the dm6 blacklist. Genes within each class were sorted in descending order of total signal and plotted using plotHeatmap.

### RNA extraction

Whole RNA from *OregonR*, *mute^1281^* null, or *mute^1281^* heterozygous sibling were extracted from three biological replicates of 50 stage 14 *Drosophila* embryos. Embryos were handpicked into embryo wash buffer (120mM NaCl, 0.07% Triton-X100), dechorionated as described above, and washed with ddH_2_O to remove detergents. Embryos were homogenized in 25ul of ice cold TRIzol (Invitrogen 15596018) with a pestle then the final volume was brought up to 300ul total TRIzol. Samples were incubated at RT for 5 min, spun down for 1 minute at 15,000 xg and the supernatant containing RNA was removed to a fresh tube.

RNA was then purified using the Zymo Direct-zol RNA miniprep kit (R2050) and quantified using the QUBIT RNA HS Assay Kit (Thermo, Q32852). We found the DNase1 step included in the Zymo kit was not enough to fully remove histone genomic DNA, so the purified RNA was then digested with dsDNase (Thermo EN0771) before any subsequent use.

### First strand synthesis and Reverse Transcriptase (RT)-qPCR

First strand cDNA synthesis was performed using the SuperScript III Reverse transcriptase kit (Thermo, 18080044) with random hexamers (Thermo, N8080127) to amplify non-polyadenylated histone RNAs. No RNAseH digestion was performed. Samples were then used for qPCR analysis with Luna® Universal qPCR Master Mix (NEB M3003). Data represented is the average of three technical replicates per primer set of three biological replicates (nine total) for each genotype which was normalized to α-tubulin-84D. ΔΔC^t^ was calculated using *OregonR* as the control for the other two genotypes. Primers used were:

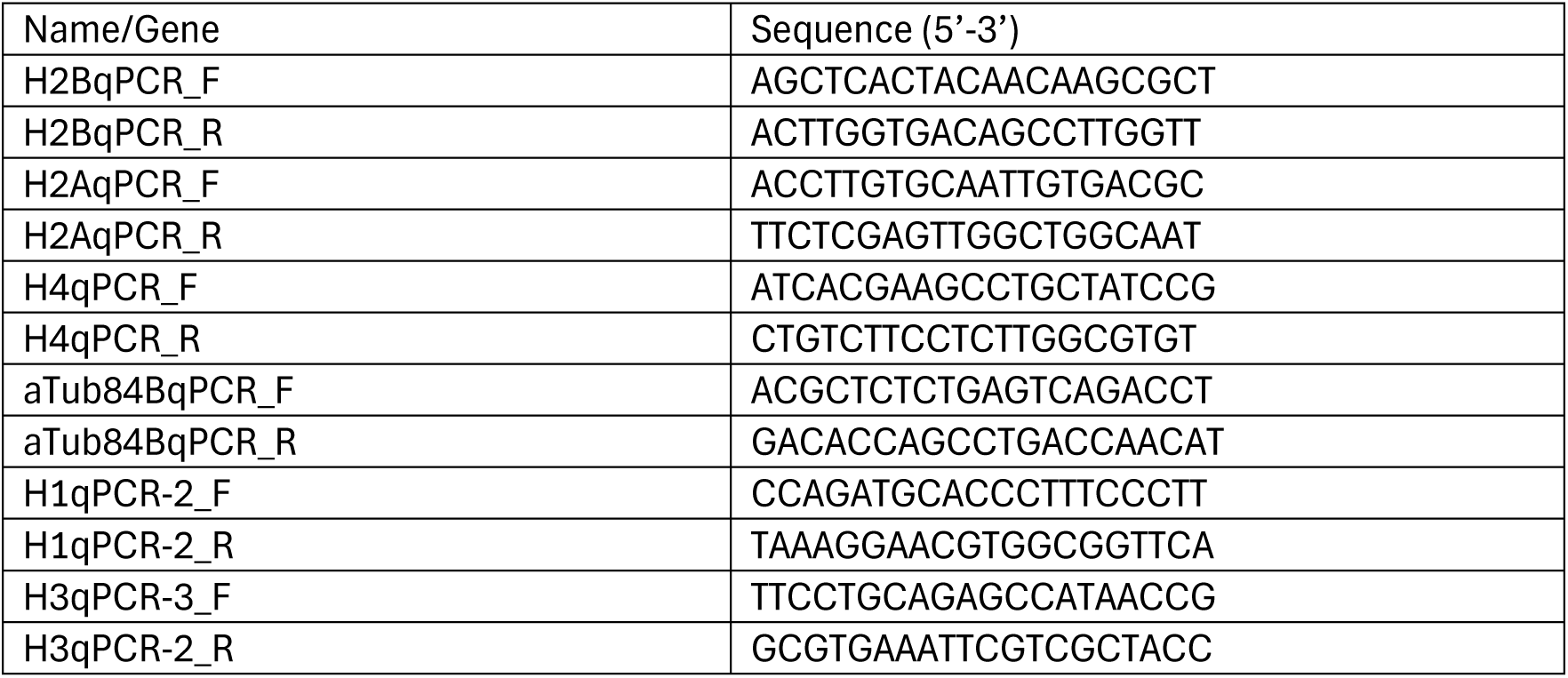

### RNA-sequencing library preparation

RNA-seq libraries were prepared using the Universal Plus total RNA-seq with NuQuant (Tecan) as described in (Crain, Butler, et al., 2024). Pooled libraries were sequenced PE50 at the UNC High Throughput Sequencing Facility (HTSF) on a NextSeq2000 P1 flow cell.

### RNA-seq data processing and analysis

Three replicates for each genotype were processed using the nf-core/rnaseq pipeline (v3.22.2; doi: 10.5281/zenodo.1400710) (Ewels et al., 2020) executed via Nextflow (v24.04.2) using the Apptainer/Singularity container profile (unc_longleaf). Raw FASTQ files were trimmed and filtered using Trim Galore! (v0.6.7). Read quality was assessed by FastQC before and after trimming, and summary reports were aggregated using MultiQC (v1.33). Trimmed reads were aligned to to the custom dm6-chrHis assembly using the STAR aligner with Salmon (v1.10.1) quantification. Aligned reads were coordinate-sorted and indexed using SAMtools (v1.21) and duplicate reads flagged using Picard MarkDuplicates.

Gene-level transcript abundance was quantified using Salmon in alignment-based mode against the STAR-aligned BAMs. Salmon was also run independently in mapping-based (decoy-aware) mode directly from the trimmed FASTQ reads against a transcriptome index built from the genome FASTA and GTF as recommended. Transcript-level Salmon estimates were aggregated to gene-level count matrices using tximport, producing the merged gene count matrix and gene length normalization matrix, used as input for differential abundance analysis. RNA-sequencing Differential gene expression analysis was performed using the nf-core/differentialabundance pipeline (version 1.5.0; doi: 10.5281/zenodo.7568000) which utilizes DEseq2 (Love et al., 2014) to assess differential gene expression between genotypes. Gene Set Enrichment Analysis (GSEA) was performed on the DEseq2 output using PANGEA (PAthway, Network and Gene-set Enrichment Analysis), a multi-species enrichment tool (Hu et al., 2025). Gene sets used were Direct Gene Ontology (GO) Biological Processes (with p< 0.01) and preferred tissue, built using *Drosophila* modEncode RNA-seq. Volcano plot of differential gene expression (Log2FC) vs significance (-log10 adj. p-val) was generated in R (v4.5.0) with ggplot2.

### Comparison of Mute Cut&Run signal versus differential gene expression

To assess Mute signal at differentially expressed genes and all RD-histone genes, Per-gene signal statistics were extracted from each compressed matrix file (from deepTools computeMatrix) and mergerd with RNA-seq differential expression results using a custom Python script. Volcano plot of differential gene expression (Log2FC) vs Summed Mute signal (CPM) was generated in R (v4.5.0) with ggplot2.

### Wing disc twin spot analysis

To generate a timecourse of differently aged control and Mute null mosaic tissues in a wildtype background in 3^rd^ instar larval wing imaginal discs, yw,hsFLP;FRT42D, UbiGFP/Cyo flies were mated with either yw,hsFLP;FRT42D, UbiRFP flies (control), or yw,hsFLP;FRT42D,Mute^1281^/Cyo-twi-GFP flies (Mutant) and let lay in vials for 48 hours at 25°C. The flies were then flipped out and the vials were aged at 25°C and heatshocked at 37°C for 10 minutes either 72, 60, 48, or 24 hours prior to dissection (72, 84, 96, or 120hrs AED) at wandering 3^rd^ instar (144 hrs AED). Larvae were genotyped by the presence of Ubi-GFP (and Ubi-RFP in control) and the lack of Cyo-twi-GFP signal to select either yw,hsFLP; FRT42D, UbiGFP/FRT42D, UbiRFP or yw,hsFLP; FRT42D,UbiGFP/FRT42D, Mute^1281^ larvae. Once genotyped, larvae were dissected, fixed, and stained as previously described (Kemp et al., 2025)

For generating a timecourse of control and Mute null mosaic tissues in the reduced competition of a Minute (M(2)53(1)) background, yw,hsFLP;FRT42D, UbiRFP, M(2)53(1)/Cyo flies were mated with either yw,hsFLP;FRT42D, UbiGFP flies (control), or yw,hsFLP;FRT42D,Mute^1281^/Cyo-twi-GFP flies (Mutant) and let lay in vials for 48 hours at 25°C. The flies were then flipped out and the vials were aged at 25°C and heatshocked at 37°C for 10 minutes either 72, 60, 48, or 24 hours prior to dissection (120, 132, 144, or 168hrs AED) at wandering 3^rd^ instar (196 hrs AED), which was delayed due to the Minute background. Larvae were genotype as described above to select either yw,hsFLP; FRT42D, UbiRFP, M(2)53(1)/FRT42D, UbiGFP or yw,hsFLP; FRT42D,UbiRFP, M(2)53(1)/FRT42D, Mute^1281^ larvae, and dissected and stained as described above.

## Data Availability

Sequencing data and custom script will be made available upon publication

