## Supplemental Figures and Legends for "Differential control of both cell cycle-regulated and quantitative histone mRNA expression by *Drosophila* Mute"

**Supplemental Figure S1. Mute and Mxc signal at non-histone genomic bins**

**A-C)** IGV screenshots of Mute, Mxc, and IgG signal at chr2L 707kb window (A), chr Y 49kB window (B), and chr3R 25 kB and chr3L 26kB windows (C). The removed HisC Locus is highlighted with blue box in A. **D)** Manhattan plot of IgG signal (CPM) in 5080bp bins across entire *Drosophila* genome. HGC nrormalized IgG signal at ChrHis is shown by red dot, top five bins for IgG signal are labeled. **E)** Metagene plot and heatmap of summed HGC normalized IgG signal across all *Drosophila* genes +/- 0.5 kb on either side. Genes were clustered using unsupervised hierarchical clustering for n = 3 bins as in Fig 1. Histone genes located in cluster 3 (asterisk). **F)** Scatter-plot of Mute vs Mxc signal (CPM) at each 5080bp genomic bin. HGC signal at chrHis is shown by red dot, top five signal bins are labeled.

**Supplemental Figure S2. *mute* mutant ganglion mother cells inappropriately express core RD histone mRNAs**

**A)** VNC of stage 14 homozygous Mute deficiency (Df) (bottom) and heterozygous siblings (top) stained with antibodies against Mute and FLASH. Mute-LS antibody recognizes both short and long isoform of Mute. Scale bar = 5μm **B)** Schematic of Type I neuronal lineage in the Drosophila VNC. **C)** VNC of *OregonR* and *mute*$^{1281}$ null embryos stained with antibodies recognizing the GMC marker Prospero (cyan) and the neuroblast marker Deadpan (magenta) and hybridized with the CDS probe (yellow, shown in panel D). White boxes indicate regions shown in panel C. Scale bar = 20μm. **D)** Higher magnification view of images from panel B. White dotted circles indicate GMCs expressing core RD histone mRNA in the *mute* mutant but not wild type. Yellow dotted circle indicates cell with high levels of prospero, deadpan and histone mRNA. Scale bar = 2μm. **E-G)** Stacked bar graphs of quantified co-occurrence of Deadpan and Prospero (E), Prospero and histone RNA (F), and Deadpan and Histone RNA (G) in *OreR* and *mute*$^{1281}$ embryos as seen in C. Significance determined by unpaired t-test with Welch correction and FDR. Error bars = SD ; n = 4; ** = pval <0.005; * = pval < 0.05.

**Supplemental Figure S3. RNA-FISH of individual histone RNAs show Mute controls temporal and quantitative output of histone gene expression.**

**A)** Schematic of a single RD histone gene unit and the location of multiple fluorescently labeled oligonucleotide probe sets used to detect transcription and accumulation of core RD histone mRNAs including CDS (pink), H1 (yellow), H2a (green), and H3 (blue). **B)** Log10 transformed histogram of mean H2a and CDS probe set in each genotype as seen in Fig 4D,E. Solid line represents mean of n=5 with SEM shaded bars. Dashed vertical lines represent average 90$^{th}$ percentile start for each genotype. **C)** Histograms of pixel intensity distributions for H3, H1, H2a, and CDS probe sets in each genotype. Solid line represents mean of n=5 with SEM shaded bars. Dashed vertical lines represent average median for each genotype. **D)** Bar graph of median pixel value for each probe set across genotypes. * = padj < 0.05; Error bars = SD; n = 5 . **E)** Bar graph of Log2 fold change (Log2FC) in expression for all five RD-histones measured by RT-qPCR ($\Delta\Delta C^t$) in *mute*$^{1281}$ heterozygous sibling embryos compared to *OregonR*. No significant change in expression is seen. Error bars = lfcSE; n = 3 **F)** Bar graph of Log2FC in expression for all five Replication independent histone variants measured by RNA-Seq (RPKM) in *mute*$^{1281}$ embryos compared to *OregonR*. No significant change in expression is seen. Error bars = lfcSE; n = 3.

**Supplemental Figure S4. Differentially expressed genes in *mute*$^{1281}$ embryos enriched in muscle processes, tissues.**

**A,B)** IGV browser shot of RPKM normalized signal of combined strands at down regulated *Ldh* (A) and up-regulated *Mef2* (B) in *OregonR* (blue) and *mute*$^{1281}$ (green) stage 14 embryos. **C)** Metagene plot and heatmap of summed HGC normalized IgG signal across all RD-histone genes (5, dark blue), upregulated genes (448, light blue) and downregulated genes (350,yellow) normalized to 2kb +/- 0.5 kb on either side. **D,E)** Bar chart of Log2FC (y-axis) and significance (color) of gene sets most enriched with DEGs from *mute*$^{1281}$ embryos. Gene sets generated by PANGEA include preferred tissue (modEncode RNA-seq) (D) and Direct Biological Process from Gene Ontology (E).

**Supplemental Figure S5. *mute* mutant wing imaginal discs cells do not proliferate even with reduced competition from neighboring cells**

**A)** Schematic of mitotic recombination using a *Minute* chromosome. **B)** Schematic of expected twin spots if reduced competition in the *Minute* background allows *mute*$^{1281}$ cells to proliferate and not be eliminated from the wing disc epithelium. **C)** Twin spots resulting from mitotic recombination induced at the indicated times before dissection using a control wild type chromosome. Scale bar = 10μm. **D)** Twin spots resulting from mitotic recombination induced at the indicated times before dissection using a *mute*$^{1281}$ null mutant chromosome. Note that *mute* mutant clones do not appear even in the *Minute* background. Scale bar = 10μm.

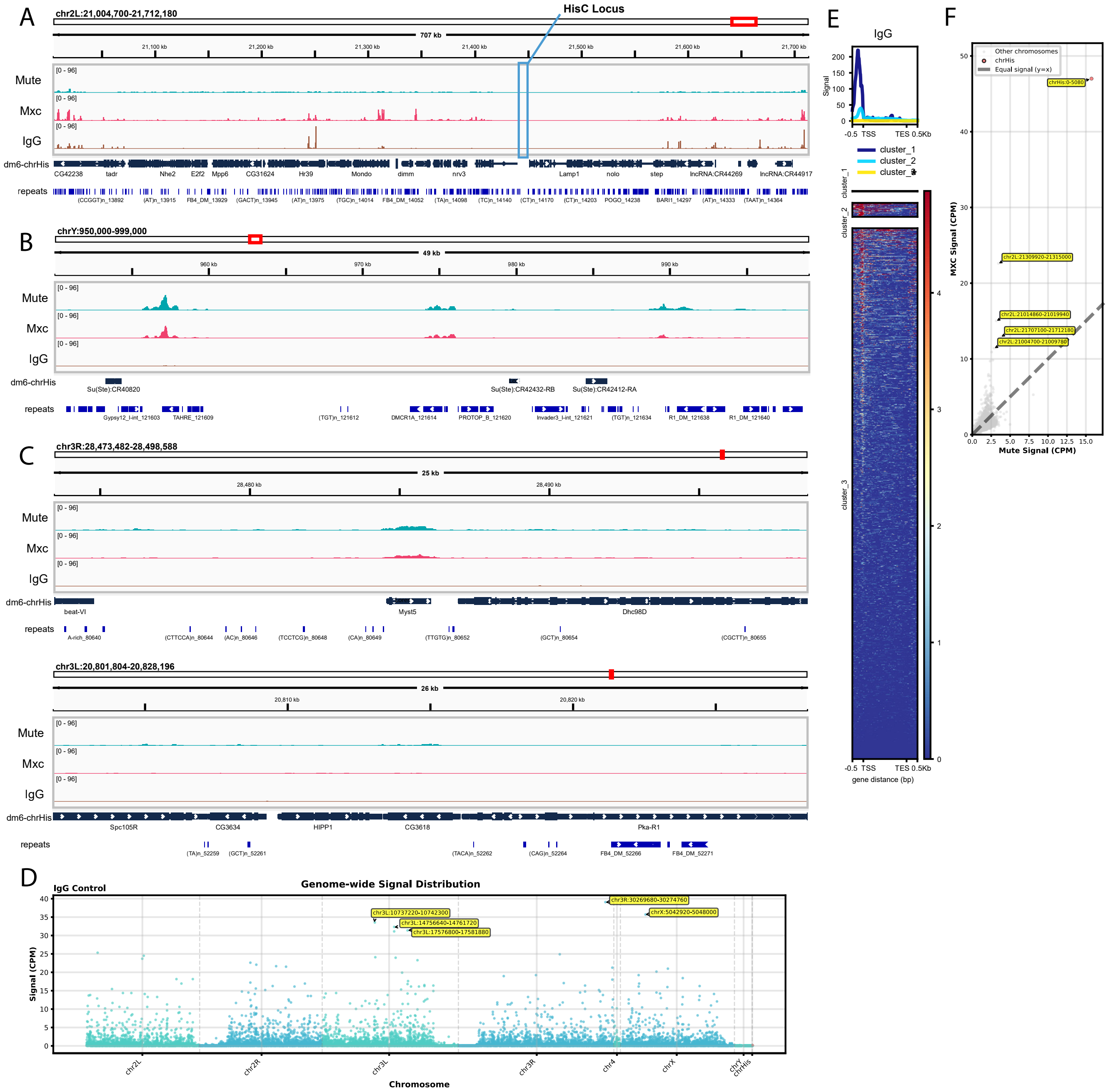
A
chr2L:21,004,700-21,712,180
HisC Locus
707 kb
21,100 kb
21,200 kb
21,300 kb
21,400 kb
21,500 kb
21,600 kb
21,700 kb
Mute
Mxc
IgG
[0 - 96]
dm6-chrHis
CG42238
tadr
Nhe2
E2f2
Mpp6
CG31624
Hr39
Mondo
dimm
nrv3
Lamp1
nolo
step
lncRNA:CR44269
lncRNA:CR44917
repeats
B
chrY:950,000-999,000
49 kb
960 kb
970 kb
980 kb
990 kb
Su(Ste):CR40820
Su(Ste):CR42432-RB
Su(Ste):CR42412-RA
C
chr3R:28,473,482-28,498,588
25 kb
28,480 kb
28,490 kb
beat-VI
Myst5
Dhc98D
chr3L:20,801,804-20,828,196
26 kb
20,810 kb
20,820 kb
Spc105R
CG3634
HIPP1
CG3618
Pka-R1
D
IgG Control
Genome-wide Signal Distribution
Signal (CPM)
Chromosome
chr3R:30269680-30274760
chrX:5042920-5048000
chr3L:10737220-10742300
chr3L:14756640-14761720
chr3L:17576800-17581880
chr2L
chr2R
chr3L
chr3R
chr4
chrX
chrY
chrHis
E
IgG
Signal
-0.5 TSS
TES 0.5Kb
cluster_1
cluster_2
cluster_3
gene distance (bp)
F
Other chromosomes
chrHis
Equal signal (y=x)
chrHis:0-5080
chr2L:21309920-21315000
chr2L:21014860-21019940
chr2L:21707100-21712180
chr2L:21004700-21009780
MXC Signal (CPM)
Mute Signal (CPM)

Suppl Fig S1

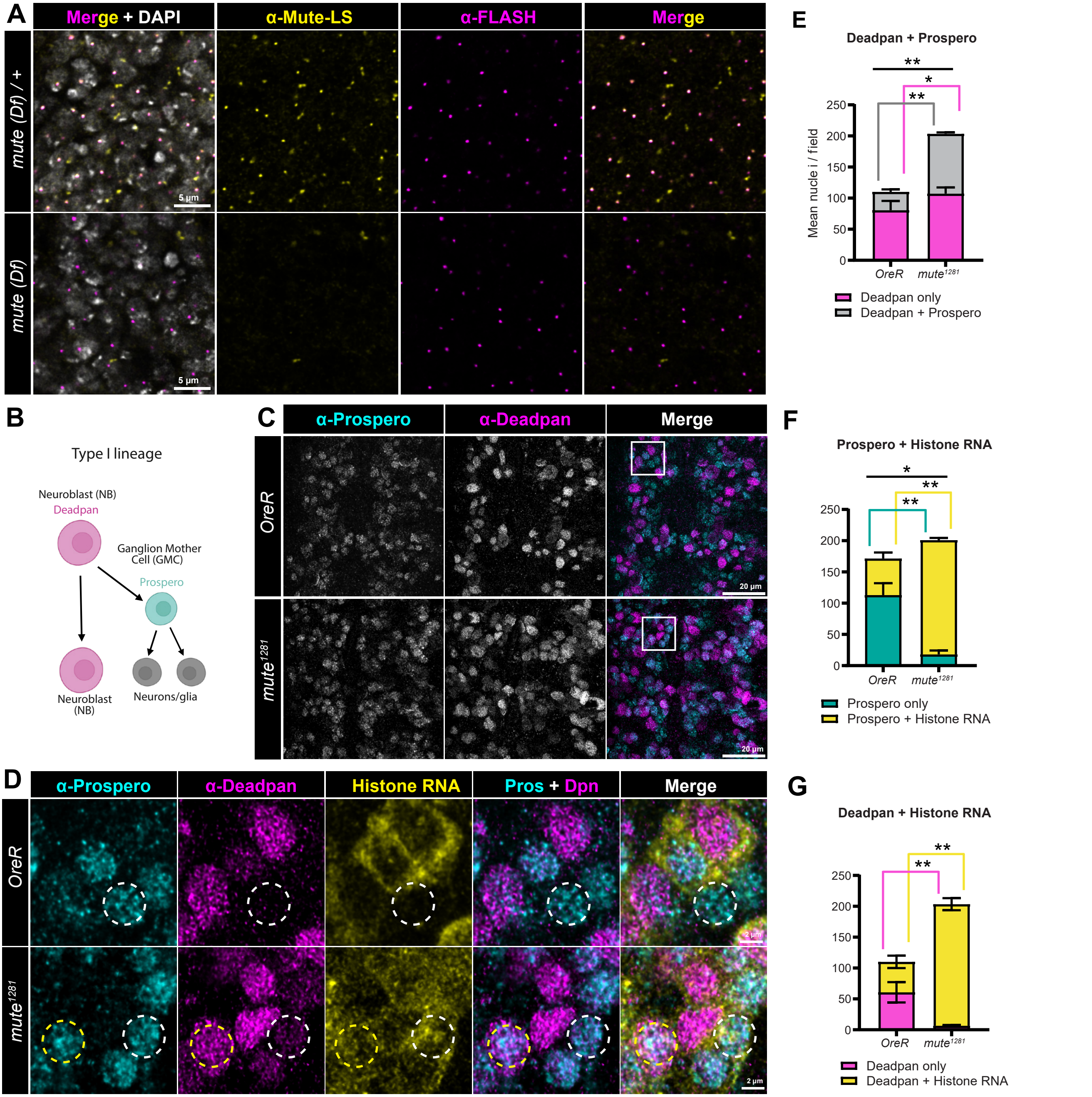
A
Merge + DAPI
α-Mute-LS
α-FLASH
Merge
mute (Df) / +
mute (Df)
5 µm
5 µm
E
Deadpan + Prospero
Mean nucle i / field
OreR
mute1281
Deadpan only
Deadpan + Prospero
B
Type I lineage
Neuroblast (NB)
Deadpan
Ganglion Mother Cell (GMC)
Prospero
Neuroblast (NB)
Neurons/glia
C
α-Prospero
α-Deadpan
Merge
OreR
mute1281
20 µm
20 µm
F
Prospero + Histone RNA
OreR
mute1281
Prospero only
Prospero + Histone RNA
D
α-Prospero
α-Deadpan
Histone RNA
Pros + Dpn
Merge
OreR
mute1281
2 µm
2 µm
G
Deadpan + Histone RNA
OreR
mute1281
Deadpan only
Deadpan + Histone RNA

Suppl Fig S2

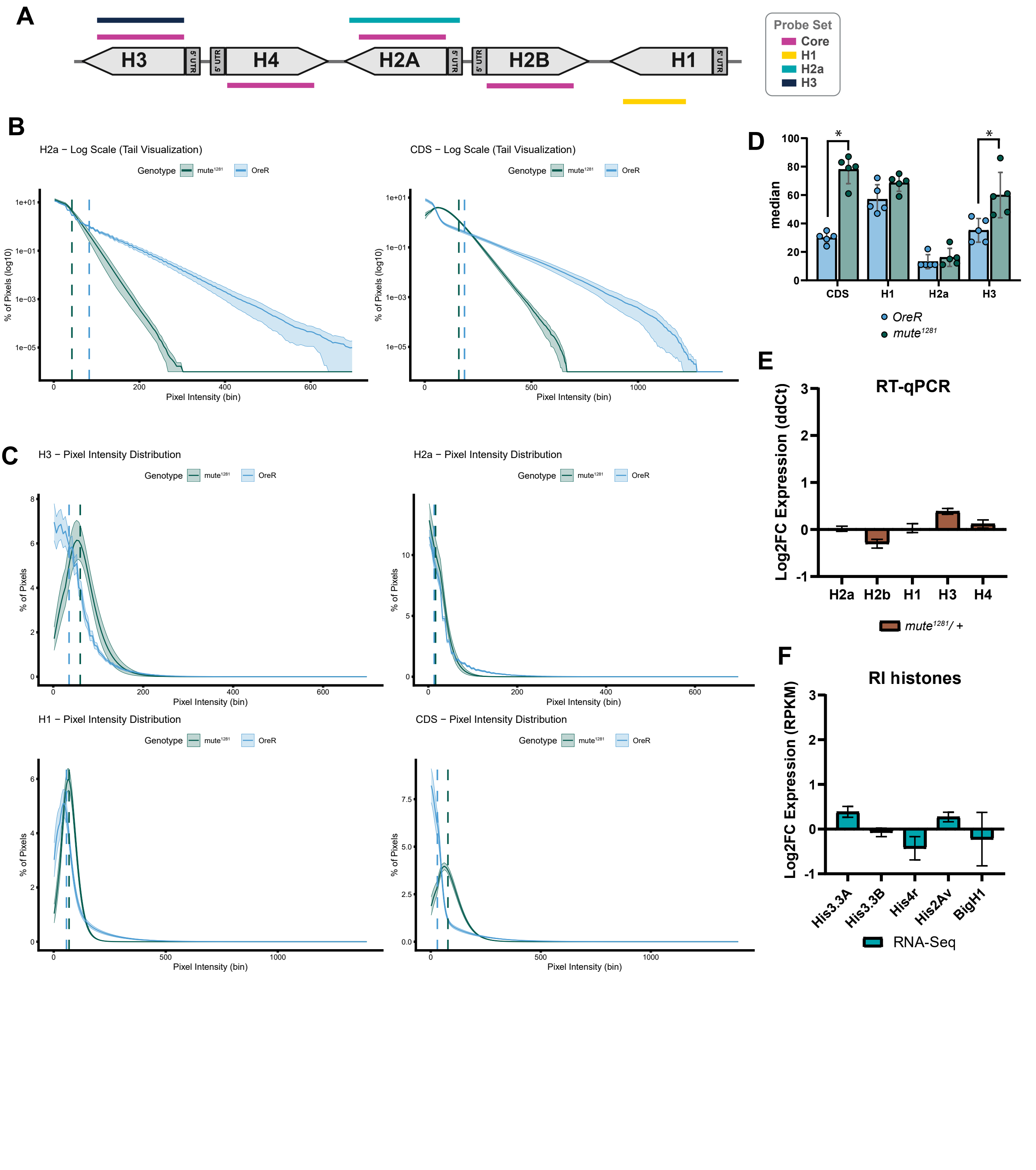

A
Probe Set
Core
H1
H2a
H3
H3
5' UTR
5' UTR
H4
H2A
5' UTR
5' UTR
H2B
H1
5' UTR
B
H2a – Log Scale (Tail Visualization)
Genotype
mute1281
OreR
% of Pixels (log10)
Pixel Intensity (bin)
CDS – Log Scale (Tail Visualization)
D
median
CDS
H1
H2a
H3
OreR
mute1281
E
RT-qPCR
Log2FC Expression (ddCt)
H2a
H2b
H1
H3
H4
mute1281/ +
C
H3 – Pixel Intensity Distribution
% of Pixels
H2a – Pixel Intensity Distribution
H1 – Pixel Intensity Distribution
CDS – Pixel Intensity Distribution
F
RI histones
Log2FC Expression (RPKM)
His3.3A
His3.3B
His4r
His2Av
BigH1
RNA-Seq


Suppl Fig S3

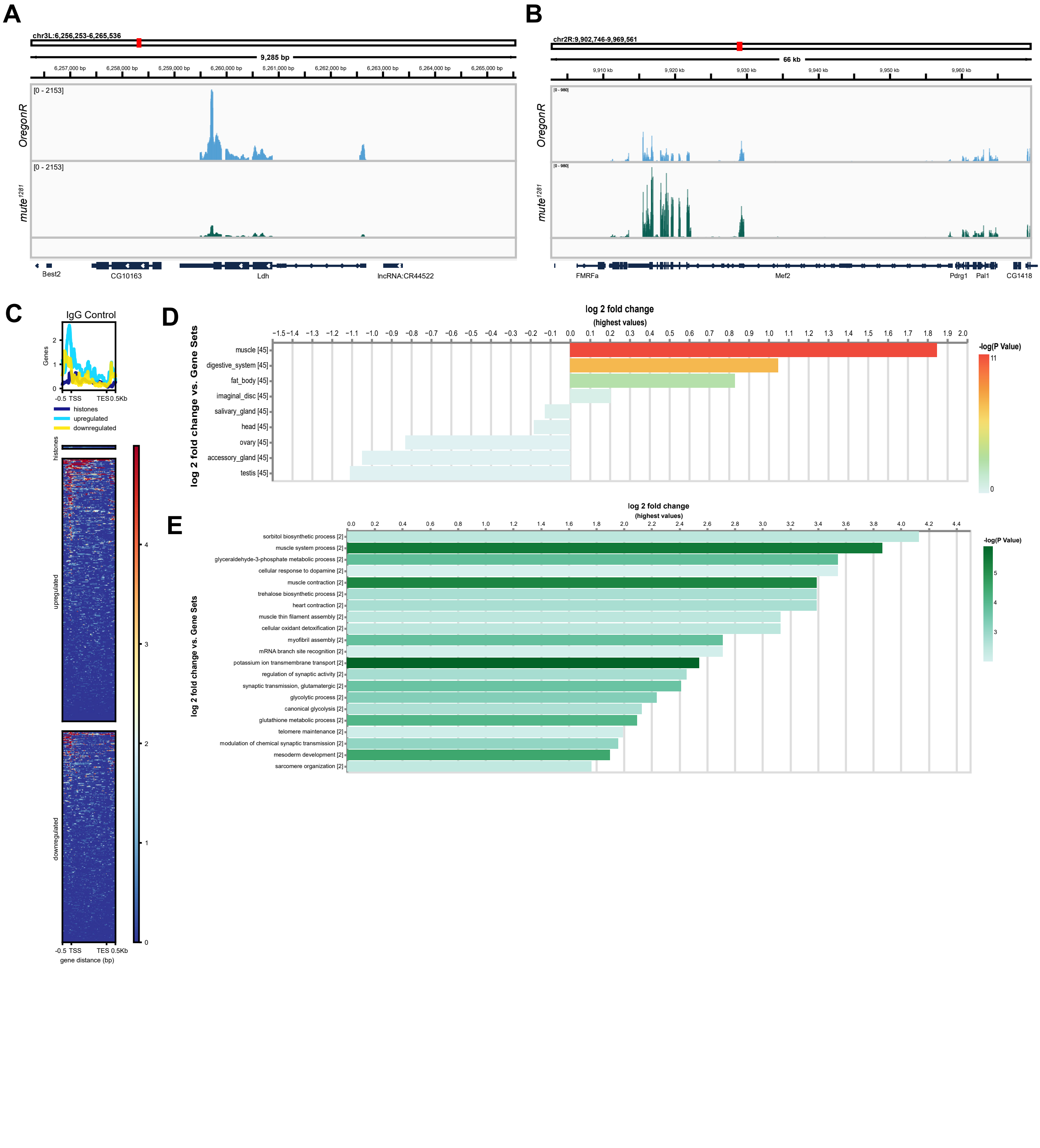

A
chr3L:6,256,253-6,265,536
9,285 bp
OregonR
mute1281
[0 - 2153]
Best2
CG10163
Ldh
lncRNA:CR44522
B
chr2R:9,902,746-9,969,561
66 kb
FMRFa
Mef2
Pdrg1
Pal1
CG1418
C
IgG Control
histones
upregulated
downregulated
-0.5 TSS
TES 0.5Kb
gene distance (bp)
D
log 2 fold change
(highest values)
log 2 fold change vs. Gene Sets
-log(P Value)
muscle [45]
digestive_system [45]
fat_body [45]
imaginal_disc [45]
salivary_gland [45]
head [45]
ovary [45]
accessory_gland [45]
testis [45]
E
log 2 fold change
(highest values)
log 2 fold change vs. Gene Sets
-log(P Value)
sorbitol biosynthetic process [2]
muscle system process [2]
glyceraldehyde-3-phosphate metabolic process [2]
cellular response to dopamine [2]
muscle contraction [2]
trehalose biosynthetic process [2]
heart contraction [2]
muscle thin filament assembly [2]
cellular oxidant detoxification [2]
myofibril assembly [2]
mRNA branch site recognition [2]
potassium ion transmembrane transport [2]
regulation of synaptic activity [2]
synaptic transmission, glutamatergic [2]
glycolytic process [2]
canonical glycolysis [2]
glutathione metabolic process [2]
telomere maintenance [2]
modulation of chemical synaptic transmission [2]
mesoderm development [2]
sarcomere organization [2]


Suppl Fig S4

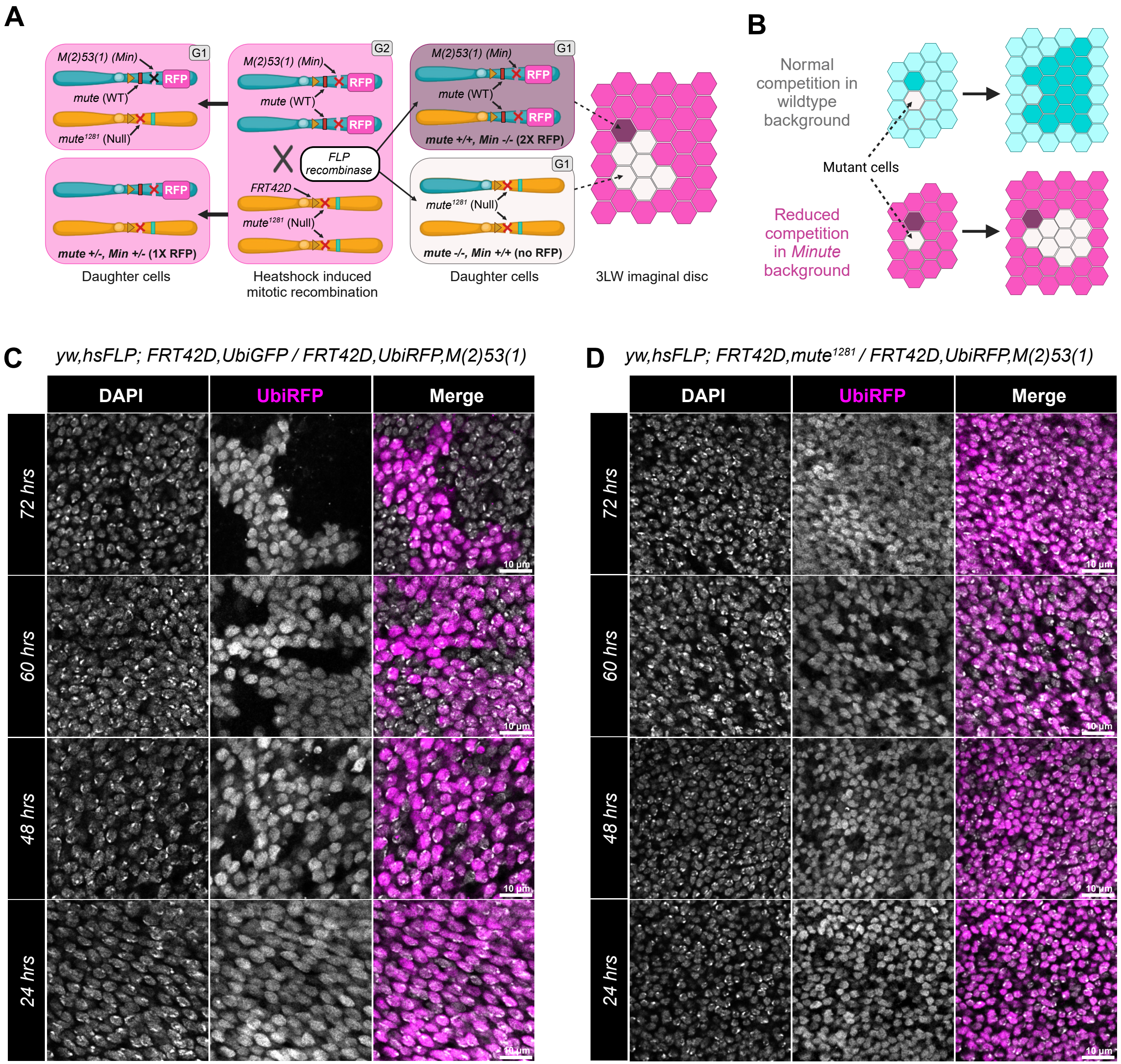
A
M(2)53(1) (Min)
G1
RFP
mute (WT)
mute1281 (Null)
G2
M(2)53(1) (Min)
RFP
mute (WT)
RFP
FLP
recombinase
FRT42D
mute1281 (Null)
G1
M(2)53(1) (Min)
RFP
mute (WT)
RFP
mute +/+, Min -/- (2X RFP)
G1
mute1281 (Null)
mute -/-, Min +/+ (no RFP)
RFP
mute +/-, Min +/- (1X RFP)
Daughter cells
Heatshock induced
mitotic recombination
Daughter cells
3LW imaginal disc
B
Normal
competition in
wildtype
background
Mutant cells
Reduced
competition
in Minute
background
C
yw,hsFLP; FRT42D,UbiGFP / FRT42D,UbiRFP,M(2)53(1)
DAPI
UbiRFP
Merge
72 hrs
60 hrs
48 hrs
24 hrs
10 µm
D
yw,hsFLP; FRT42D,mute1281 / FRT42D,UbiRFP,M(2)53(1)
DAPI
UbiRFP
Merge
72 hrs
60 hrs
48 hrs
24 hrs
10 µm

Suppl Fig S5
